# Whole genome duplication potentiates inter-specific hybridisation and niche shifts in Australian burrowing frogs *Neobatrachus*

**DOI:** 10.1101/593699

**Authors:** Polina Yu. Novikova, Ian G. Brennan, William Booker, Michael Mahony, Paul Doughty, Alan R. Lemmon, Emily Moriarty Lemmon, Levi Yant, Yves Van de Peer, J. Scott Keogh, Stephen C. Donnellan

## Abstract

Polyploidy has played an important role in evolution across the tree of life but it is still unclear how polyploid lineages may persist after their initial formation. While both common and well-studied in plants, polyploidy is rare in animals and generally less well-understood. The Australian burrowing frog genus *Neobatrachus* is comprised of six diploid and three polyploid species and offers a powerful animal polyploid model system. We generated exome-capture sequence data from 87 individuals representing all nine species of *Neobatrachus* to investigate species-level relationships, the origin and inheritance mode of polyploid species, and the population genomic effects of polyploidy on genus-wide demography. We resolve the phylogenetic relationships among *Neobatrachus* species and provide further support that the three polyploid species have independent autotetraploid origins. We document higher genetic diversity in tetraploids, resulting from widespread gene flow specifically between the tetraploids, asymmetric inter-ploidy gene flow directed from sympatric diploids to tetraploids, and current isolation of diploid species from each other. We also constructed models of ecologically suitable areas for each species to investigate the impact of climate variation on frogs with differing ploidy levels. These models suggest substantial change in suitable areas compared to past climate, which in turn corresponds to population genomic estimates of demographic histories. We propose that *Neobatrachus* diploids may be suffering the early genomic impacts of climate-induced habitat loss, while tetraploids appear to be avoiding this fate, possibly due to widespread gene flow into tetraploid lineages specifically. Finally, we demonstrate that *Neobatrachus* is an attractive model to study the effects of ploidy on the evolution of adaptation in animals.

## Introduction

Polyploidy or whole genome duplications (WGDs) play important roles in ecology and evolution (1, 2). Although polyploidization predominantly occurs in plants, polyploidy has also played an important role in animal evolution. For instance, two ancient WGDs occurred early in the vertebrate lineage (3), while more recent WGDs occurred in several animal groups, including insects, molluscs, crustaceans, fishes, amphibians and reptiles (4–6). While the majority of polyploid animals switch to different atypical modes of bisexual reproduction after polyploid formation (4, 7, 8), amphibians, and more specifically anurans, are unique among polyploid animals in exhibiting multiple independent occurrences of diploid and sexually reproducing polyploid sister species (9).

While polyploidization has thus occurred frequently across the tree of life, the evolutionary benefits of WGDs remain elusive. Although polyploids of hybrid origin (allopolyploids) may benefit from heterosis due to increased genetic variation, instantaneous shifts into intermediate or new ecological niches, and the redundancy of independently segregating gene copies (2, 10), newly formed allopolyploids are simultaneously subject to several disadvantages perhaps most prominently of which is their low abundance compared to the established non-polyploids (10). Conversely, while autopolyploids retain many of the same disadvantages as allopolyploids, the advantages of autopolyploidy are much less clear (11).

Here, we focus on a group of widely distributed, endemic Australian burrowing frogs, *Neobatrachus.* This genus comprises six diploid *(N. albipes, N. fulvus, N. pelobatoides, N. pictus, N. sutor, N. wilsmorei;* 2n=24) and three tetraploid *(N. aquilonius, N. kunapalari, N. sudellae;* 4n=48) species (12, 13), all characterised by bisexual reproduction (14). Tetraploid species of *Neobatrachus* were suggested to have independent origins based on mitochondrial DNA (mtDNA) (15) and to have originated through autotetraploidy rather than allotetraploidy, as they exhibit tetrasomic inheritance and show high rates of tetravalent formation during meiosis (14, 16). The diploid species are well defined based on external morphology, male advertisement calls and divergence at allozyme loci (17–19). Frog call structure differs among ploidies with higher ploidy species generally having lower pulse rates, a trait linked to nuclear volume increase with increasing ploidy (20). Tetraploid *Neobatrachus* species have lower pulse number and rate in their advertisement calls compared to diploids with multiple pulses in their calls (however *N. sutor* (2n) and *N. wilsmorei* (2n) have calls with a single pulse), and each of the *Neobatrachus* species retain distinct calls (21, 22). This differs from the more extensively studied gray treefrog, *Hyla versicolor*, where tetraploids have also originated from multiple independent origins, but have similar calls, and so have merged into a single species through frequent interbreeding (23).

It has also been observed that, while the *Neobatrachus* tetraploid species are distributed sympatrically with some of the diploid species, they are also able to occupy more arid areas across Australia (2, 15). Generally, polyploidy has been associated with greater tolerance to harsher conditions, but it is not clear whether whole-genome duplication broadly provides a fitness advantage or simply is a consequence of elevated rates of unreduced gamete formation (2, 10, 24, 25), which might be more prone to occur in extreme environments. At the same time, changing environments may contribute to amphibian extinction rates, which continue to increase and are driven by many interdependent factors such as habitat loss, emergence and spread of diseases, invasive species and pollution (26–29).

In the current study, we use an anchored hybrid enrichment approach (AHE) (30–33) to resolve the phylogenetic relationships among *Neobatrachus* species, and to assess fine-scale intra-specific genetic population structure. We also quantify the extent of hybridization between the nine *Neobatrachus* species with a particular focus on taxa with contrasting ploidies. Finally, we combine population dynamics assessments with changes in ecologically suitable areas for each species to describe population responses to climate changes.

## Results

### Phylogenetic relationships, genetic population structure and gene flow in *Neobatrachus*

We first generated sequence data and alignments for 439 targeted orthologous nuclear loci of 87 *Neobatrachus* individuals spanning the entire genus as well as nine *Heleioporus* individuals as outgroups (see Methods). We then built a species tree and gene trees from the sequenced loci with ASTRAL-II (34) using RaxML (35) (Fig. 1A). These traditional bifurcating species-tree methods reveal extensive conflict among genealogies, as well as between nuclear DNA (nDNA) and mtDNA topologies, but they do all demonstrate that the three tetraploid species do not form a monophyletic group (Fig.1A, Fig. S1, Fig. S2). Multidimensional scaling (MDS) of gene tree topologies suggested that nuclear loci constitute either two or four topology clusters (Fig. S3), indicating competing signal. In the absence of informative fossil material, we estimated the approximate evolutionary timescale of *Neobatrachus* frogs for mitochondrial (Fig. 1C) loci using secondary calibrations (36). Interspecific divergence times provided support for an Eocene-Oligocene origin of *Neobatrachus* (Fig. 1C). Clustering patterns of gene tree topologies, together with relatively deep divergence times between the species, suggest recent or ongoing gene flow. Although similar patterns could be expected for allopolyploid (hybrid) origins of the tetraploids, based on modeling allele frequency distributions for allo- and autotetraploids, our results seem to support the previously suggested autotetraploid origin of *Neobatrachus* tetraploid species (Fig. S4, see Methods) (14, 15).

**Fig. 1.**
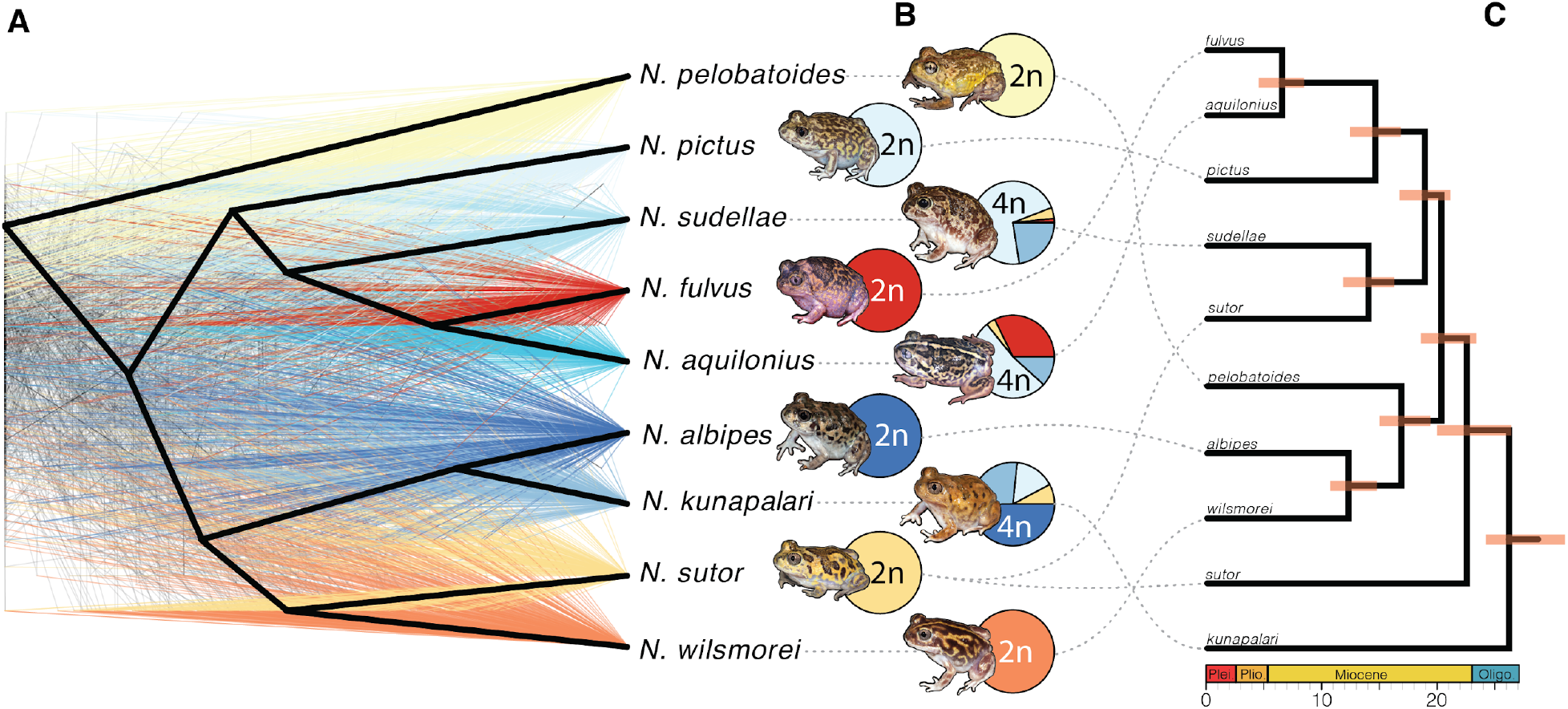
Independent origins of *Neobatrachus* tetraploids and high levels of reticulation. (A) Gene trees, colored by clade, for 361 nuclear loci based on 2 individuals per species show considerable incongruence and differ from the nuclear (bold black topology in (A) and mitochondrial (C) species trees. Conflict between gene tree clusters (Fig. S3), the nuclear species tree, and the mitochondrial tree suggest non-bifurcating relationships between the species. (B) Pie charts represent summarised admixture proportions for each species (summing assignments for each individual, Fig. S1, Fig. 2) at optimal clustering with K=7. Tetraploids *(N. sudellae, N. aquilonius* and *N. kunapalari)* show highly admixed ancestries. (C) Dated mitochondrial tree built from coding sequences conflicts with the topology of the nuclear species tree in A. Red bars represent 95% confidence intervals on the ages of nodes, noted in millions of years before present.

To investigate the population structure of *Neobatrachus* further we assessed it by ADMIXTURE analyses (37) on extracted polymorphism data (66,789 sites in total; see *Methods;* Fig. 1B, Fig. 2 Fig. S1, Fig. S5). Overall, admixture clustering corresponded with the phylogenetic placement of the individuals on the species tree (Fig. 1, Fig. S1). Diploid species were clearly split at K=7 and did not show admixed individuals, whereas all three tetraploid species showed admixture with each other and with local diploid species (Fig. 1, Fig. 2, Fig. S1).

**Fig. 2.**
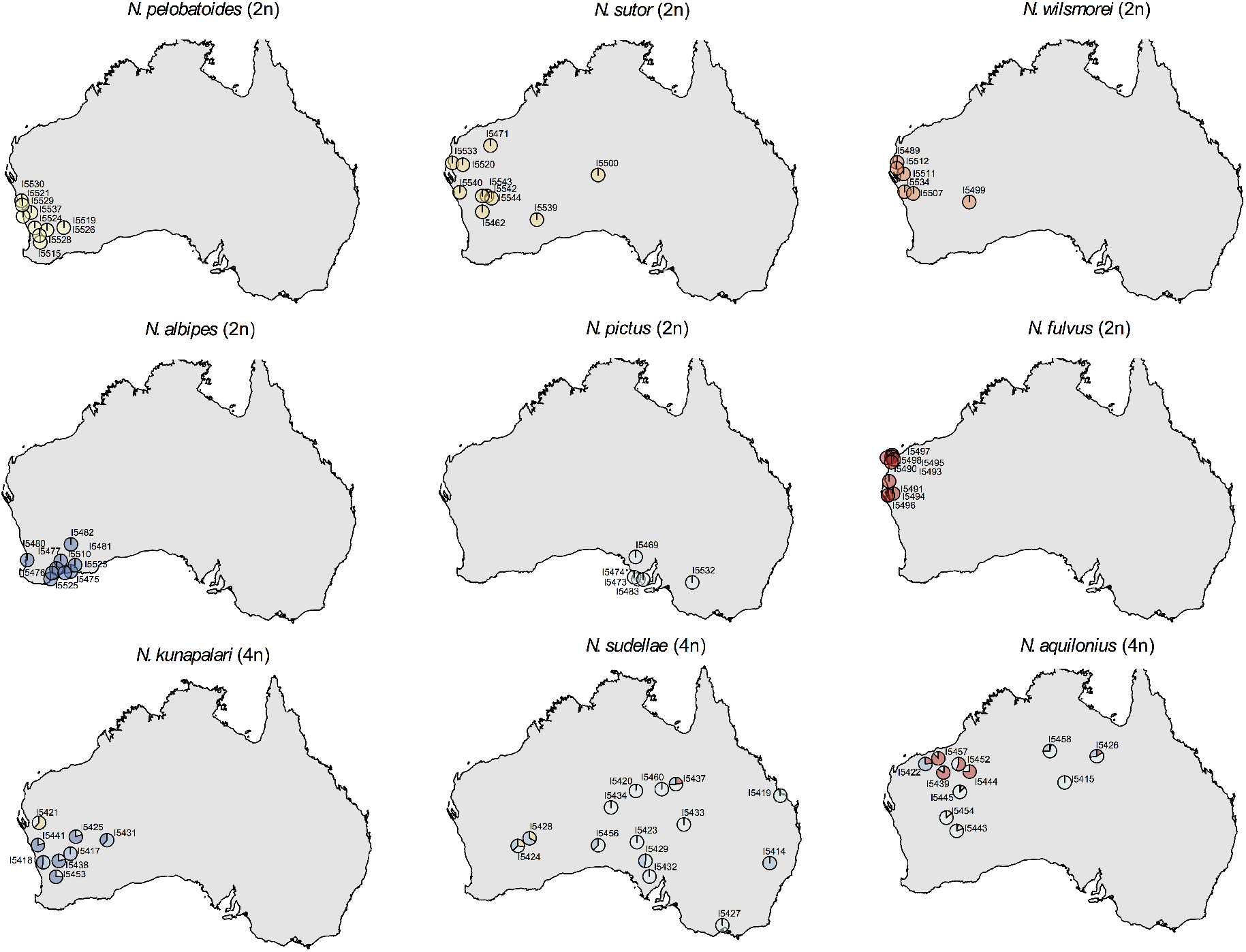
ADMIXTURE results (K=7) shown separately for each species. According to the geographical locations of the sampled individuals, pie charts show the probability of the assignment of the individual to one of the 7 individually colored clusters. Overlapping pie charts on the map have been moved just enough to appear separate. Diploid *Neobatrachus* species (top 6: *N. pelobatoides*, *N. albipes*, *N. wilsmorei, N. sutor*, *N. pictus*, *N. fulvus)* are each assigned to separate clusters, while all three tetraploid species (bottom 3: *N. kunapalari, N. sudellae*, *N. aquilonius)* show inter-species admixture.

To further assess the complex demographic history of *Neobatrachus*, we performed TreeMix (38) modeling where species relationships are represented through a graph of ancestral populations (Fig. 3A). The structure of the graph is inferred from allele-frequency data and Gaussian approximation of genetic drift such that the branch lengths in the graph are proportional to the amount of drift since population split. We sequentially added up to 15 migration events, showing saturation of the model likelihood at five additional migration edges on average for 30 runs of Treemix, each with a different seed for random number generation (Fig. 3E). We show an example of the inferred introgression events and the bifurcating graph for the model with five migration events for the run that resulted in the highest maximum likelihood (Fig. 3A-D). Inferred migration events (Fig. 3B) indicate widespread directional introgression and interploidy gene flow between the polyploid species, however, only two introgression events had p-value lower than 0.05 in this particular run: from *N. sudellae* (4n) to *N. kunapalari* (4n) and from *N. sutor* (2n) to *N. kunapalari* (4n). Since there was some variability in inferred migration edges from run to run, to estimate the most frequently inferred migration events we summed the significant inferred migration edges among 30 TreeMix runs with five events allowed (Fig. 3F). Migration events were found most frequently from *N. sudellae* (4n) to *N. kunapalari* (4n) (19 of 30 runs) and from *N. sutor* (2n) to *N. kunapalari* (4n) (12 of 30 runs). Interploidy introgression events were mostly asymmetric and from diploids to tetraploids, which corresponds with our ADMIXTURE cluster assignment results (Fig. 1, Fig. 2). Inferred introgression events are broadly congruent with clusters of conflicting gene-tree topologies (Fig. S3). Each tetraploid *Neobatrachus* species *(N. aquilonius, N. kunapalari, N. sudellae)* is sister to a diploid species in the TreeMix graphs (Fig. 3A-B, tips highlighted in bold) as well as in the species trees (Fig. 1), which is consistent with previously suggested independent origins for the tetraploid species (15).

**Fig. 3.**
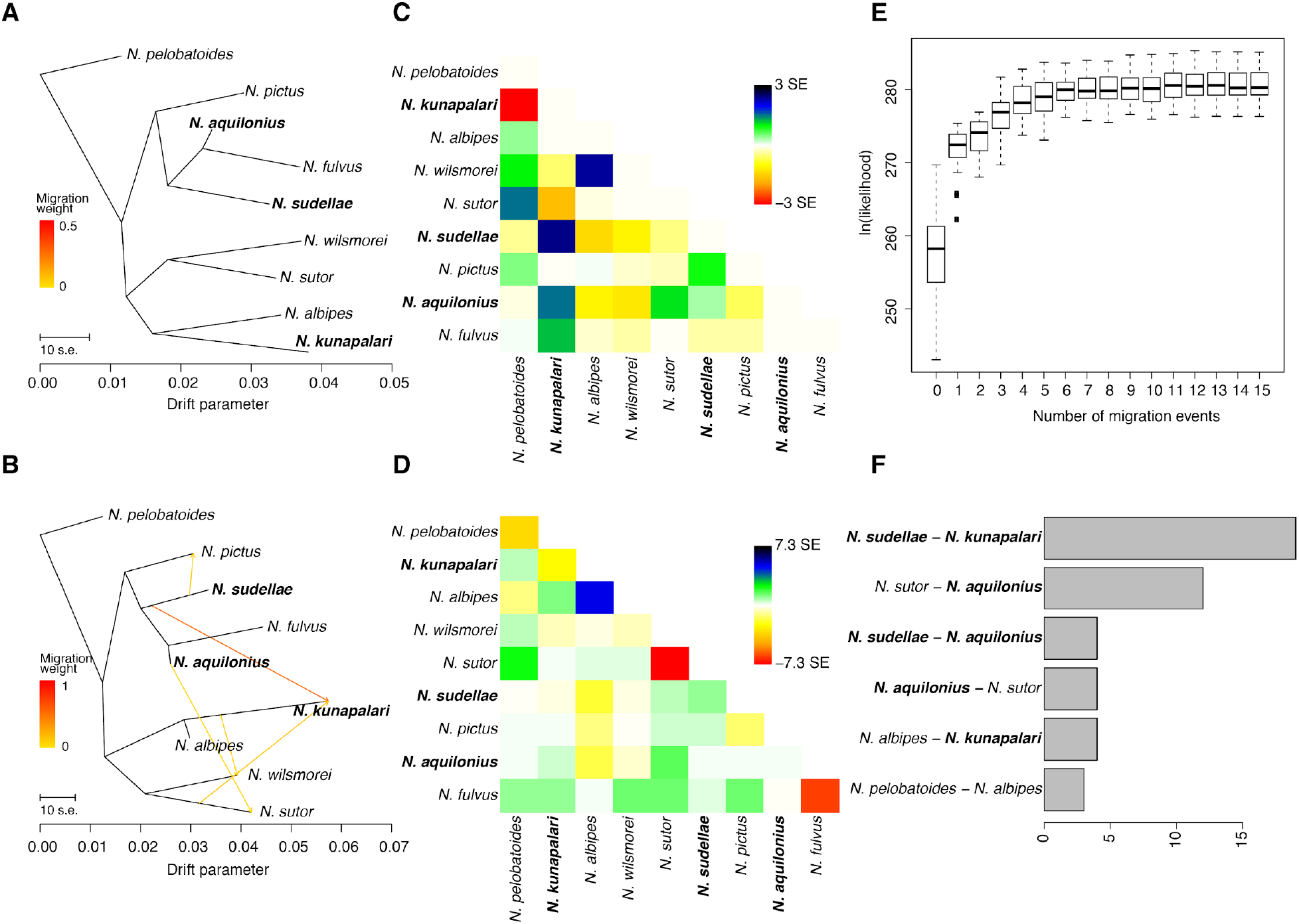
Widespread introgression between *Neobatrachus* species. (A) Bifurcating maximum likelihood tree produced by TreeMix. (B) Example of a graph produced by TreeMix with 5 allowed migration events. (C) Scaled residual fit between observed data and predicted model in (A). Plot shows half of the residual covariance between each pair of populations divided by the average standard error across all pairs. Positive residuals represent populations where the model underestimates the observed covariance, meaning that populations are more closely related to each other in the data than in the modeled tree. Such population pairs are candidates for admixture events. Similarly, negative residuals indicate pairs of populations where the model overestimates the observed covariance. Overall, the residual plot of the model suggested that model fit could be improved by additional edges (migration events). (D) Scaled residual fit between observed data and predicted model in (B). Compared to Fig. 3C this suggests that, although the complexity of the species relatedness is not fully represented by the model, major gene flow events and their direction were probably captured. (E) Box plots of 30 runs of TreeMix (each started with a different seed for random number generation) likelihood at different numbers of allowed migration events; saturation starts after 3 additional migration edges. (F) Bar plot showing the number of times a particular directional migration event was inferred in 30 TreeMix runs with 5 migration events allowed. We show only the events which were inferred more than twice.

### Estimation of suitable distribution areas and demographic patterns

Tetraploid species have the highest nucleotide diversity among *Neobatrachus* species (Fig. 4D, Supplementary Table 1), which is most likely due to gene flow directed to tetraploid taxa and introgression between tetraploids of different origin. This is supported also by Fst distances (Fig. S6), where Fst distances between tetraploid species are the lowest, while Fst distances between tetraploids and diploid are larger, and Fst distances between the diploid lineages are the highest, suggesting stronger isolation.

**Fig. 4.**
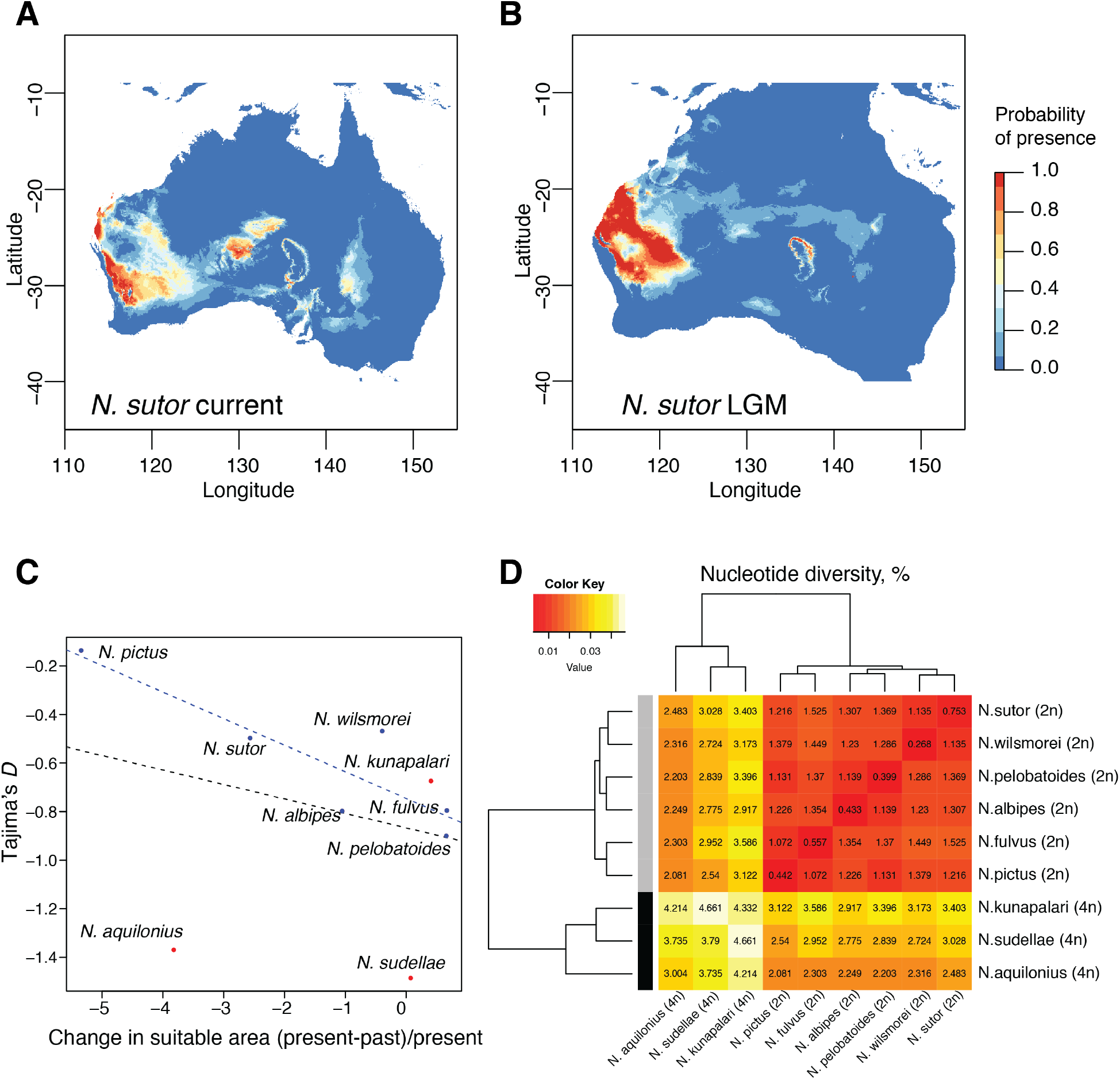
Diversity and differentiation of *Neobatrachus* species and geographical suitability estimates. (A) Example of the estimation of the suitable distribution area for *N. sutor*, based on occurrence data and current climate. (B) Example of the projection of the suitable distribution area for *N. sutor* based on the past climate at around 20Kya at LGM (last glacial maximum). (C) Scatter plot showing relative change of the predicted suitable area at the LGM and current conditions for each species as a function of Tajima’s D estimator. Diploid species show high correlation between Tajima’s *D* and distribution area change (blue line, Pearson’s correlation −0.88 (R^2^ =0.72, p-value=0.02); (D) Hierarchical clustering of *Neobatrachus* species based on mean nucleotide diversity within and between the species.

To estimate the dynamics in population abundance over recent times, we measured Tajima’s *D*, a summary statistic that measures the lack or excess of rare alleles in a population compared to the neutral model. All of the *Neobatrachus* species have negative median values of Tajima’s D, which suggests that none of the species are experiencing dramatic population diversity decline (Fig. 4B). We used the observed Tajima’s *D* values as a proxy for each species’ demographic patterns and compared them with estimated change in the suitable geographic area (Fig. 4A-C). In order to describe the ecological areas occupied by the different *Neobatrachus* species, as well as changes in those areas since the last glacial maximum (LGM) at around 20 Kya, we made use of occurrence data (39) (Fig. S7) and climate datasets (40). We first performed a PCA of reduced bioclimatic variables concentrating on one of the highly correlated variables (r>0.85, Pearson correlation coefficient; see *Methods)* for individuals from the *Neobatrachus* occurrence data (Fig. S8). Using this substantially increased geographic sampling compared to our sequenced sample set, we could see moderate clustering of the individuals by species, which demonstrates that *Neobatrachus* species differ in their ecological (climatic) occupancies.

We then modelled suitable distribution areas for each species separately with MaxEnt (41), which applies machine learning maximum entropy modeling on the climate data at the geographical locations of the species occurrence data. Bioclimatic variables had different impacts on the model for each species (Fig. S9), however, appeared to be more similar for sympatric species (for example, *N. sutor* and *N. wilsmorei*) than for allopatric species (for example, *N. pictus* compared to any other diploid species). By projecting the models built on the current climate data on the past climate data (at the last glacial maximum, LGM) we could estimate the changes in the suitable habitat area since the LGM relative to the current suitable area for different species (Fig. 4A-C, Fig. S10-11). We observed a correlation between the change in the suitable habitat area and median Tajima’s *D* for the diploid *Neobatrachus* species (Fig. 4C). As shrinkage in suitable habitat areas increases for a given diploid species, Tajima’s *D* values also increase. This suggests that climate change may have already had a negative effect on diploid *Neobatrachus’* genetic diversity via loss of suitable habitat even if populations are not obviously already getting smaller. Interestingly, tetraploid species appear to be the outliers to this trend, which we suggest may be due to their highly admixed genetic structure.

## Discussion

We have investigated the population genomic consequences of genome duplication in a vertebrate model by generating and analysing nucleotide sequence data for 439 loci in 87 *Neobatrachus* individuals, covering the entire genus, including the three currently recognized tetraploids. The observation of non-bifurcating relationships between closely related species (Fig. 1) is now common (42–52), and is well understood to be caused by either incomplete lineage sorting or gene flow, or both. Population structure analysis revealed that each of the diploid species forms discrete clusters, consistent with their status as phylogenetically distant species. The tetraploids, however, were assigned to a mixed set of clusters, suggesting gene flow between each other and with the diploid species from overlapping geographical areas (Fig. 1, 2; Fig. S1). Extended analysis of the population structure and potential gene flow uncovered migration (gene flow) events from *N. sudellae* (4n) to *N. kunapalari* (4n) and from *N. sutor* (2n) to *N. aquilonius* (4n). In several analyses, we inferred migration from *N. sudellae* (4n) to *N. aquilonius* (4n), reversed migration between *N. sutor* and *N. aquilonius*, migration from *N. albipes* (2n) to *N. kunapalari* (4n) and even *N. pelobatoides* (2n) to *N. albipes* (2n). The latter, if true, may be attributed to an ancient migration event, since we do not see any evidence for recent mixing between the diploids.

Unidirectional gene flow from diploid to tetraploids could be envisioned through a couple of scenarios. Firstly, a diploid individual could produce an unreduced gamete in a cross with a tetraploid forming a tetraploid that could backcross with the sympatric tetraploid species. While cytological evidence of the production of unreduced gametes in diploids is available in the form of triploid hybrids between diploid *Neobatrachus* species, we presently do not have direct evidence of this first scenario for inter-ploidy gene flow in *Neobatrachus* (Appendix 1, Supplementary Table 3, Fig. S12) (18, 22, 53–55). Secondly, a diploid individual could cross with a tetraploid and form a triploid. A triploid producing an unreduced gamete (3n) in a cross with a diploid (normal 1n gamete) would produce a tetraploid that could backcross to a tetraploid individual producing introgression from the diploid into the tetraploid gene pool, i.e. the so-called “triploid bridge” (56). We have cytological and molecular genetic evidence of naturally occurring triploid *Neobatrachus* and direct cytological evidence of triploid *Neobatrachus* producing balanced 3n gametes (Appendix 1). In another polyploid frog complex, *Odontophrynus*, triploids produced in laboratory crosses formed balanced haploid, diploid and triploid gametes (53), which opens the possibility of triploids crossing with diploids or tetraploids to produce tetraploids that could backcross into the sympatric tetraploid gene pool. Selective mating scenarios based on call characteristics could increase the likelihood of the backcross matings with tetraploids (54). It is also worth mentioning that *Neobatrachus* species are explosive breeders and require very heavy rains (>25 mm in a rainfall) to promote emergence. In such events, sympatric species, for example *N. kunapalari, N. pelobatoides* and *N. albides* can all be breeding in the same ponds or pools (Dale Roberts, personal communication). A recent study on *Bufo japonica*, also an explosive breeder, showed that multiple paternity occurred commonly in that species in ponds with high densities of amplexed pairs but not in ponds with low density populations (57). The inferred cause was stray sperm in the pond from other amplexed pairs or possibly unpaired males.

A mechanism for unreduced gamete formation, i.e. temperature stress, has been identified experimentally (24, 25). Similar brief stress conditions during zygote development before the first cell division could block the first division and form a tetraploid that could cross into a sympatric tetraploid gene pool (58). Cold or heat shocks are possible for *Neobatrachus* egg clutches at numerous locations across southern Australia that experience temperature extremes of the order of −5 °C to 48 °C. Breeding by more southerly *Neobatrachus* tends to follow the onset of autumn and winter rains, typical of the Mediterranean climate in this region. Exposure to freezing or near freezing conditions in shallow egg deposition sites following cold fronts associated with rain fall events is frequently possible. *Neobatrachus* in the arid zone breed in association with heavy summer rains, which may be associated with heat wave conditions.

Extensive gene flow between *Neobatrachus* species, especially between the tetraploid species, makes it difficult to estimate the true ancestral diploid population(s) for the tetraploids. Previously, it has been suggested that tetraploid *Neobatrachus* species might have independent origins (15). Our results place tetraploid species in a polyphyletic arrangement on the species tree and on the TreeMix graphs, and suggest at least two independent autopolyploid origins: genetically closest diploid lineages to *N. aquilonius* and *N. sudellae* are *N. fulvus* and *N. pictus*, while the closest diploid lineage to *N. kunapalari* is *N. albipes.* An important question that remains is what allows admixture between *Neobatrachus* tetraploids of potentially different origin and admixture of tetraploids with the local diploids, while the diploids seem to be currently isolated from each other? It may be that gene flow for close diploid relatives into tetraploids occurs frequently enough subsequent to the formation of the tetraploids to maintain enough phenotypic similarities for continued mating opportunities. Similarly, the tetraploid tree frogs *Hyla versicolor* of multiple origins show high levels of interbreeding, however the levels of divergence between the ancestral diploid species in this case might be shallower (23). Another example of a similar pattern was shown in plants, where polyploidy is more frequent: diploid *Arabidopsis lyrata* and *A. arenosa* could not hybridize, while tetraploidy seems to overcome the endosperm-based hybridization barrier enabling gene flow between the two species (59, 60).

*Neobatrachus* species are widely distributed in Australia with tetraploid species occurring more in the central (drier) area compared to diploids, which is reflected on the principal component analysis of the climatic data for species occurrences (Fig. S8). Areas occupied by different *Neobatrachus* species differ only slightly in their environmental characteristics (Fig. 8). Worth mentioning is that climatic variables do not entirely describe ecological niches, which could differ in other characteristics such as timing of breeding and foraging, food source preference, etc. Nevertheless, ecological niche modelling based on climate data may provide additional insights into population dynamic trends. Here, we applied the MaxEnt modelling approach to the publicly available climate and occurrence data for all nine *Neobatrachus* species, comparing the present and past suitable geographical areas. Most of the *Neobatrachus* species showed substantial changes of the suitable areas comparing current and past presence probabilities (Fig. S10-11). Interestingly, the estimated change in suitable habitat areas and population genetics estimator of demographic trends (Tajima’s D), obtained from independent datasets, were correlated (Fig. 4C). Tetraploid species appear to be outliers from the general trend, probably due to their mixed population structure: in this case, emergence of rare alleles in the population due to migration events will affect Tajima’s *D* estimator. Overall, it appears that the species with greater shrinkage of suitable area since the last glacial maximum had less negative median Tajima’s *D* values, which suggests an ongoing shift from population expansion to population contraction.

## Conclusion and Outlook

*Neobatrachus* frogs represent a group of diploid and tetraploid species with a complex ancestry. Our results, revealing gene flow between tetraploids and asymmetric inter-ploidy gene flow, pose a number of important questions concerning the evolution of sexual polyploid animals. Whole-genome sequencing data for *Neobatrachus* species would not only help to refine the population structure and introgressive mixing in the genus, but also provide information on potential adaptive effect of the introgressed regions. One could hypothesize that a wide and potentially rapid spread of the tetraploids into new territories was facilitated by introgression from the locally adapted diploids (61), and that more detailed sampling of the tetraploids from parts of their distribution ranges remote from sympatry with diploids may reveal evidence of early versus ongoing gene flow. Moreover, population-level genomic resequencing of multiple diploid and tetraploid sister species could provide insight into the unique biology of autotetraploid sexual animals and effects of the tetraploidization on their evolution. The results on the changing suitable habitat areas for the *Neobatrachus* species highlight the importance of continuous observation of their population dynamics. Monitoring the current status of biodiversity through collection of species occurrence data and population genetic data allows the prediction of population dynamics and hopefully timely response in conservation efforts in the face of rapidly changing environments (62, 63). Emerging methods of public engagement to collect occurrence and other data (video and audio; www.frogid.net.au, (64, 65)) have potential to provide essential information on the state of frog species.

## Methods

### Anchored Hybrid Enrichment (AHE) phylogenomics

All the samples examined were obtained from the Australian Biological Tissue Collection at the South Australian Museum. Details of all samples examined are presented in the Supplementary Data. We collected AHE data at Florida State University’s Center for Anchored Phylogenomics (www.anchoredphylogeny.com), following the methods described in Lemmon et al. (30) and Prum et al. (66). Briefly, after quantifying the extracted DNA using Qubit, we sonicated the DNA to a size range of 150-500bp using a Covaris Ultrasonicator. We then prepared indexed libraries using a Beckman Coulter FXp liquid-handling robot. After library QC using Qubit, we pooled the libraries in groups of 16 and enriched the library pools using an hybrid enrichment kit developed for use in Anurans (31, 32). Finally, we sequenced the enriched library pools on two lanes of an Illumina 2500 sequencer with a PE150 protocol at the Translational Laboratory at Florida State University.

Following sequencing, we quality filtered the reads using the Casava high-chastity filter, then demultiplexed the reads using the 8bp indexes with no mismatches tolerated. To increase read length and correct for sequencing errors, we merged read pairs that overlapped by at least 18bp using the method of Rokyta et al. (67). This process also removed sequencing adapters. We then performed a quasi ‘de novo’ assembly of the reads following Hamilton et al. (68), with *Pseudacris nigrita*, and *Gastrophryne carolinensis* as references. In order to reduce the potential effects of low level sample contamination, we retained only the assembly clusters containing more than 61 reads. In order to produce phased haplotypes from the assembly clusters, we applied the Bayesian approach developed by Pyron et al. (69), in which reads overlapping polymorphic sites are used to identify the likely phase of allelic variants within each locus. Because this approach was developed to accommodate any ploidy level, we were able to isolate two or four haplotypes for diploid and tetraploid individuals, respectively. We determined orthology for each locus using a neighbor-joining approach based on pairwise sequence distances, as described in Hamilton et al. (68). We aligned homologous haplotypes using MAFFT v7.023b (70), then auto-trimmed/masked the alignments following the approach of Hamilton et al. (68), but with *MINGOODSITES=12, MINPROPSAME=0.3*, and *MISSINGALLOWED=48.* Final alignments were visually inspected in Geneious R9 (Biomatters Ltd., (71)) to ensure that gappy regions were removed and misaligned sequences were masked.

### Whole mitochondrial genomes

Mitochondrial bycatch from exome capture methods can be an appreciable source of mitogenomic data (72). To take advantage of these data, we assembled mitochondrial genomes from raw reads using custom scripts built around existing assembly and alignment software. Raw reads for each sample were assembled against the mitochondrial genome of *Lechriodus melanopyga* (73), using a baiting and mapping approach implemented in MITObim (74). Mapped reads from each individual genomes were then aligned using MUSCLE (75), and adjusted manually. Entire concatenated mitogenomes were used to investigate sample identities, and to build an initial phylogenetic tree. To estimate divergence dates between species using mitochondrial data, we extracted 13 protein coding regions (CDS), and partitioned them separately. We then trimmed the concatenated mitogenome phylogeny and CDS alignments down to two representatives of each taxon except for *Neobatrachus sutor*, which was not monophyletic (and therefore was trimmed down to four samples), and included two species of *Heleioporus* and *Lechriodus* as outgroups. The resulting tree and alignments were used as inputs for baseml and ultimately mcmctree, which we ran until we reached 20,000 post-burnin samples.

### Phylogenetic analysis

To generate a molecular species tree, we started by reconstructing individual genealogies for each of the 439 recovered loci. We analyzed two datasets, of which the first included all samples (except triploids discussed in the Results section), while the second was trimmed down to just two individuals per species. Results from the full sampling can be found in the *Supplementary Material* (Fig. S2), and the finer sampling in the main text (Fig. 1). We used RaxML (35) to simultaneously search for the best tree and apply 100 rapid bootstraps, implementing the GTRGAMMA model of nucleotide evolution for each locus. In generating species trees, coalescent methods have been shown to be more accurate than concatenation in case of extensive incomplete lineage sorting, and so we used the shortcut coalescent method ASTRAL III. Shortcut coalescent methods like ASTRAL take individual gene trees as input, and are much more computationally efficient than full coalescent analyses. We used our RAxML-generated gene trees as input for ASTRAL, allowing us to make use of all our molecular data.

To address gene-tree incongruence and investigate possible conflicting signals in our data, we used multidimensional scaling (MDS) to approximate the relative distances between gene tree topologies (76). To prepare the data, we trimmed down gene trees to one sample per species of *Neobatrachus*, and discarded loci missing any taxa, leaving us with 361 loci. We started by simply visualizing gene-tree incongruence overlaying the topologies of all 361 loci in DensiTree (Fig. 1). We then calculated the pairwise distances between all gene trees using the Robinson-Foulds metric, in the R package APE (77). We projected the tree distances into two and three dimensions (representing tree topology space) using MDS, as visualizing and interpreting more dimensions is difficult. To test if gene trees are uniformly distributed throughout tree space, or clustered, we used the partitioning around the medoids algorithm as implemented in the R package CLUSTER(78). We chose the optimum number of clusters (k), using the gap statistic, calculated for each *k* = 1–10. Clusters of gene trees represent similar topologies, and so we then summarized each cluster using ASTRAL, to identify consistent differences in topology.

We estimated the evolutionary timescale of *Neobatrachus* frogs for the mitochondrial DNAm, using 13 protein coding sequences, and a concatenated RAxML phylogeny of those loci as input for mcmctree. Because no valuable fossil information for *Neobatrachus* is available, we used four secondary calibrations from the most extensive anuran time-tree to date (36). We applied these secondary calibrations as truncated Cauchy distributions with 2.5% above and below the designated bounds, to the splits between (i) Myobatrachidae and Microhylidae (lower=120, upper=140), *Kalophrynus* and remaining microhylids (lower=55, upper=66), *Cophixalus* and *Liophryne* (lower=17, upper=24), and *Microhyla* and *Kaloula* (lower=40, upper=55). This analysis produced an estimate of the divergence between *Neobatrachus* and *Heleioporus* with a 95% CI of 25-45 Mya, which was used to inform the mcmctree analysis of mtDNA coding sequences. We ran analyses until we reached 20,000 post-burnin samples.

### Population structure

Maximum likelihoods of individual ancestries were estimated with ADMIXTURE (37) for 66,789 biallelic sites combining all 439 loci and allowing for maximum 20 out of 87 *Neobatrachus* individuals (excluding the outgroup species) to have missing data at each site. In order to include tetraploid samples in ancestry assignment we randomly chose two alleles for each site. We also applied minor allele frequency threshold of two percent. Ancestral population assignment showed three local minima of cross-validation errors at K equals 3, 7 and 9 (Fig. S6), with K=7 being the lowest, which we chose for the subsequent analysis as the optimal solution.

### Misidentifications in the dataset

Both phylogenetic and admixture assignments suggested that several individuals had been misidentified in the field, which is expected for morphologically similar species and in particular for some of the diploid-tetraploid species pairs, e.g. *N. fulvus* and *N. aquilonius* (17). Field sampling can be accompanied by a certain level of honest mistakes in species identification, especially for sympatric species. However, a high level of incompletely sorted polymorphisms in recently split lineages or recent hybridization events could also result in uncertain positioning of an individual. We carefully curated the dataset and made a decision to rename some of the misidentified samples or completely exclude them from the analyses based on the amount of the missing data in the assembly, ploidy estimations from the sequencing data, identification from mitochondrial sequences and on the clear placement in a different clade. Below we describe our workflow for manual curation of the dataset to exclude or rename uncertain individuals without compromising too much on the potentially real shared variation.

The multiple sequence alignment resulting from the AHE workflow contains different amounts of informative sequence and gaps for each individual. First, we calculated the informative sequence fraction (no gaps) for each individual compared to the multiple sequence alignment length and applied a threshold of at least 0.2 of informative fraction for each individual to qualify for the subsequent analysis. Based on these criteria we excluded 6 samples (Supplementary Data).

Second, we estimated the ploidy of each sample using the nQuire software (79) on the next generation sequencing data mapped to one of the outgroup species *Heleioporus australiacus* (I5549) AHE assembly as a reference. As a preparation step for nQuire, we mapped reads to the reference using the BWA-MEM algorithm from BWA (80) (version 0.7.17), used Samtools (81) (version 1.6) to sort and index the mapping and removed potential duplicates from the PCR amplification step of library preparation with picard-tools (http://broadinstitute.github.io/picard/). We used the denoised input of base frequencies generated with default parameters for the Gaussian Mixture Model utilized in nQuire to estimate ploidy levels on the basis of frequency distributions at biallelic sites. The resulted estimations can be found in Supplementary Data. We excluded 5 samples placed in a different clade compared to the rest of the samples in the corresponding lineage, where ploidy estimation confirmed their misidentification. Finally, we renamed 2 samples to a different species name with which it clustered, in cases when initial ploidy and estimated ploidy corresponded to each other (Supplementary Data).

We have also excluded from further analysis sample I5442, initially identified as *N. kunapalari*, which was estimated to be a triploid and showed high levels of admixture between a diploid *N. wilsmorei* and potentially *N. kunapalari* or *N. sudellae.* In fact, triploid individuals are known in natural populations of *Neobatrachus* (19, 21, 22, 82, 83) (Appendix 1), and could provide an explanation for the gene flow between species of different ploidy through a “triploid bridge”, when a 3n individual formed in a cross between 2n and 4n individuals and could produce a balanced haploid, diploid or triploid gametes and in the two later cases cross to a diploid and produce 4n individuals that can backcross into the sympatric 4n population (56, 84, 85).

### Inheritance mode

In order to check the inheritance mode of the three tetraploid *Neobatrachus* species we compared allele frequencies distrubutions at biallelic sites within each individual for each species with modeled expected distributions for auto and allo tetraploids. Autotetraploids were modelled combining bam files within each diploid species from the mapping bescribed above; allotetraploids were modelled combining bam files between each diploid species (Fig. S4A). Base frequencies distributions were produced using denoised algorithm from nQuire (79). As it was previously shown (86, 87) allotetraploids with disomic inheritance mode are expected to have an excess of intermediate frequency alleles, which is supported by our models as well (Fig. S4A). Performing Wilcoxon tests for the ratios between intermediate (40-60%) and rare (<30%) allele frequencies we rejected allotetraploid origin for *N. sudellae* and *N. aquilonius. N. kunapalari* showed intermediate distributions, suggesting mixed chromosomal inheritance. A mixed chromosomal inheritance pattern can be explained under several scenarios including being newly formed allopolyploid hybrids of close relation, the presence of continued gene-flow with diploids or other autopolyploid species, or the process of diploidization in an autopolyploid with tetrasomic inheritance (88). Other analyses in this study suggest extensive gene flow between *N kunapalari* and diploids as well as tetraploids (Fig. 2, Fig. 3B,F). In addition, the mitochondrial phylogenetic tree and our hierarchical clustering of nucleotide diversity analysis both support *N. kunapalari* as the oldest tetraploid lineage in this group (Fig. 1B, Fig. 4D). While we cannot rule out the possibility that *N. kunapalari* was initially formed from the hybridization of two closely related lineages, we believe extensive gene flow and the older lineage age of *N. kunapalari* is a sufficient and more likely cause for its elevated intermediate allele frequencies as compared to the other tetraploids.

### Introgression inference

The graphs representing ancestral bifurcations and migration events were produced using Treemix V.1.12 (38). Input data contained 5092 biallelic sites called at 439 loci among 9 *Neobatrachus* species with at least 20% of the data to be present at each species at each site. Position of the root was set to *N. pelobatoides* as the nuclear species tree suggested (Fig. 1A, black). To account for linkage disequilibrium we grouped SNPs in windows of size 10 using -k flag. We also generated bootstrap replicates using -bootstrap flag and subsequently allowed up to 15 migration events with flag -m. We ran TreeMix software with 30 different random number generated seeds. For graph and residuals visualisation we used R script plotting_funcs.R from the Treemix package.

### Summary statistics and demographic tendencies

We calculated summary statistics (Fig. 4A-C) with the R package “PopGenome” (89) for all the loci with more than 100 aligned sites (leaving only non-variable and biallelic sites) separately for each species for within-species statistics (nucleotide diversity, Tajima’s D, Supplementary Table 1) and in a pairwise mode for between-species statistics (Fst).

### Species distribution modelling

Bioclimatic variables were obtained from worldclim project (40) with 2.5 minutes resolution for reconstructed climate data at Last Glacial Maximum around 20Kya, averaged conditions across 1960-1990 and the most recent available conditions averaged across 1970-2000. Software DIVA-GIS 7.5 was used to trim the data to area longitude from 110 to 155 and latitude from −40 to −9. Bioclimatic variables were excluded if they were highly correlated (r>0.85, Pearson correlation coefficient) in all 3 climatic data sets, leaving for further analysis 6 bioclimatic variables in total: BIO9 = Mean Temperature of Driest Quarter, BIO10 = Mean Temperature of Warmest Quarter, BIO12 = Annual Precipitation, BIO17 = Precipitation of Driest Quarter, BIO18 = Precipitation of Warmest Quarter, BIO19 = Precipitation of Coldest Quarter.

To model the species suitable area we used software MaxEnt (v. 3.4.1.), which predicts species distribution from climate data using the species occurrences employing machine learning technique called maximum entropy modeling (41). Here we used *Neobatrachus* species occurrence data from amphibiaweb.org (39), which includes 189 entries for *N. albipes*, 282 for *N. aquilonius*, 87 for *N. fulvus*, 588 for *N. kunapalari*, 802 for *N. pelobatoides*, 699 for *N. pictus*, 707 for *N. sudellae*, 639 for *N. sutor* and 282 for *N. wilsmorei.* We used 75% of occurrence points for each species for model training and 25% for model testing with 1,000,000 background points and 10 replicates. We have trained the model on bioclimatic variables (reduced to 6 in total, described earlier) averaged across conditions 1960-1990; and then projected that model to the same set of environmental variables from the Last Glacial Maximum. The average test AUC (area under the Receiving Operator Curve) for the replicate runs for all the species was more than 0.9 (Supplementary Table 2), indicating a high performance of the models. In order to estimate which bioclimatic variable is the most important in the models we performed a jackknife test, where model performance was estimated without a particular variable and only with this particular variable in turn (Fig. S9).

We used cloglog output format of MaxEnt, which gives an estimate between 0 and 1 of probability of presence of the species in the area. In order to determine the relative change of the suitable area we used the point-wise mean values from the 10 replicates for model predictions on current and past climate (Fig. S10-11). We extracted the suitable area for both climate conditions with R library ‘raster’ (90) with 0.8 presence probability threshold and estimated change in the suitable area relative to the current suitable area as (current-past)/current.

## Data availability

The raw genomic reads generated in this study were deposited in the NCBI SRA under the BioProject PRJNA507583. BioSample SRA number for each of the individual can be found in the Supplementary Data. Multispecies alignments of phased sequences for each locus can be found at https://bioinformatics.psb.ugent.be/gdb/Neobatrachus_AHE/

## Acknowledgments

The Australian Research Council Discovery grant DP120104146 awarded to JSK and SCD supported the sequencing work. P.Y.N acknowledges postdoctoral fellowship from The Research Foundation – Flanders (FWO), 12S9618N. This work was also supported by the European Research Council (ERC) under the European Union’s Horizon 2020 research and innovation programme [grant number ERC-StG 679056 HOTSPOT], via a grant to L.Y. We thank Sarah Catalano and Steven Myers for DNA extraction and shipping samples, Mitzy Pepper for laboratory assistance, the Western Australian Museum and Dale Roberts valuable comments on the manuscript and for some of the tissue samples and Stephen Mahony for some of the frog images. We thank Sean Holland and Michelle Kortyna at Florida State University’s Center for Anchored Phylogenomics for assistance with data collection and analysis.

## References

1. Soltis DE, Visger CJ, & Soltis PS (2014) The polyploidy revolution then…and now: Stebbins revisited. Am J Bot 101(7): 1057–1078.

2. Van de Peer Y, Mizrachi E, & Marchal K (2017) The evolutionary significance of polyploidy. Nat Rev Genet 18(7):411–424.

3. Dehal P & Boore JL (2005) Two rounds of whole genome duplication in the ancestral vertebrate. PLoS Biol 3(10):e314.

4. Otto SP & Whitton J (2000) Polyploid incidence and evolution. Annu Rev Genet 34:401–437.

5. Li Z, et al. (2018) Multiple large-scale gene and genome duplications during the evolution of hexapods. Proc Natl Acad Sci U S A 115(18):4713–4718.

6. Berthelot C, et al. (2014) The rainbow trout genome provides novel insights into evolution after whole-genome duplication in vertebrates. Nat Commun 5:3657.

7. Stenberg P & Saura A (2013) Meiosis and its deviations in polyploid animals. Cytogenet Genome Res 140(2–4):185–203.

8. Neiman M, Sharbel TF, & Schwander T (2014) Genetic causes of transitions from sexual reproduction to asexuality in plants and animals. J Evol Biol 27(7):1346–1359.

9. Mable BK, Alexandrou MA, & Taylor MI (2011) Genome duplication in amphibians and fish: an extended synthesis: Polyploidy in amphibians and fish. Journal of Zoology 284: 151–182.

10. Comai L (2005) The advantages and disadvantages of being polyploid. Nat Rev Genet 6(11):836–846.

11. Parisod C, Holderegger R, & Brochmann C (2010) Evolutionary consequences of autopolyploidy. New Phytol 186(1):5–17.

12. Roberts JD (2010) Taxonomic status of the Australian burrowing frogs Neobatrachus sudelli, N. centralis and Neoruinosus and clarification of the type specimen of N. albipes. Records of the Western Australian Museum 25: 455–458.

13. Frost DR (2016) Amphibian Species of the World: an Online Reference. Version 6.0 (Date of access). Electronic Database accessible at http://research.amnh.org/herpetology/amphibia/index.html. American Museum of Natural History, New York, USA.

14. Mahony MJ & Robinson ES (1980) Polyploidy in the australian leptodactylid frog genus Neobatrachus. Chromosoma 81(2):199–212.

15. Mable BKR, J.D. (1997) Mitochondrial DNA evolution of tetraploids in the genus Neobatrachus (Anura: Myobatrachidae). Copeia 4:680–689.

16. Schmid M, Evans BJ, & Bogart JP (2015) Polyploidy in Amphibia. Cytogenetic and Genome Research 145:315–330.

17. Mahony M & Roberts JD (1986) Two new species of desert burrowing frogs of the genus Neobatrachus (Anura:Myobatrachidae) from Western Australia. Records of the Western Australian Museum 13:155–170.

18. Roberts JD, Mahony M, Kendrick P, & Majors CM (1991) A new species of burrowing frog, Neobatrachus (Anura:Myobatrachidae), from the eastern wheatbelt of Western Australia. Records of the Western Australian Museum 15:23–32.

19. Mahony M, Donnellan SC, & Roberts JD (1996) An Electrophoretic Investigation of Relationships of Diploid and Tetraploid Species of Australian Desert Frogs Neobatrachus (Anura: Myobatrachidae). Australian Journal of Zoology 44:639–650.

20. Keller MJ & Gerhardt HC (2001) Polyploidy alters advertisement call structure in gray treefrogs. Proc Biol Sci 268(1465):341–345.

21. Roberts JD (1997) Call Evolution in Neobatrachus (Anura: Myobatrachidae): Speculations on Tetraploid Origins. Copeia 1997(4):791–801.

22. Roberts JD & Edwards D (2018) The Evolution, Physiology and Ecology of the Australian Arid-Zone Frog Fauna. On the Ecology of Australia’s Arid Zone), pp 149–180.

23. Holloway AK, Cannatella DC, Gerhardt HC, & Hillis DM (2006) Polyploids with different origins and ancestors form a single sexual polyploid species. Am Nat 167(4):E88–101.

24. Mason AS & Pires JC (2015) Unreduced gametes: meiotic mishap or evolutionary mechanism? Trends Genet 31(1):5–10.

25. Pandian TJK, R. (1998) Ploidy induction and sex control in fish. Hydrobiologia 384(1–3):167–243.

26. Stuart SN, et al. (2004) Status and trends of amphibian declines and extinctions worldwide. Science 306(5702):1783–1786.

27. Collins JP (2010) Amphibian decline and extinction: what we know and what we need to learn. Dis Aquat Organ 92(2–3):93–99.

28. Hudson MA, et al. (2016) Dynamics and genetics of a disease-driven species decline to near extinction: lessons for conservation. Sci Rep 6:30772.

29. O’Hanlon SJ, et al. (2018) Recent Asian origin of chytrid fungi causing global amphibian declines. Science 360(6389):621–627.

30. Lemmon AR, Emme SA, & Lemmon EM (2012) Anchored hybrid enrichment for massively high-throughput phylogenomics. Syst Biol 61(5):727–744.

31. Barrow LN, Lemmon AR, & Lemmon EM (2018) Targeted Sampling and Target Capture: Assessing Phylogeographic Concordance with Genome-wide Data. Syst Biol 67(6):979–996.

32. Heinicke MP, Lemmon AR, Lemmon EM, McGrath K, & Hedges SB (2018) Phylogenomic support for evolutionary relationships of New World direct-developing frogs (Anura: Terraranae). Mol Phylogenet Evol 118:145–155.

33. Yuan Z-Y, et al. (2018) Natatanuran frogs used the Indian Plate to step-stone disperse and radiate across the Indian Ocean. National Science Review:nwy092–nwy092.

34. Mirarab S & Warnow T (2015) ASTRAL-II: coalescent-based species tree estimation with many hundreds of taxa and thousands of genes. Bioinformatics 31(12):i44–52.

35. Stamatakis A (2014) RAxML version 8: a tool for phylogenetic analysis and post-analysis of large phylogenies. Bioinformatics 30(9):1312–1313.

36. Feng YJ, et al. (2017) Phylogenomics reveals rapid, simultaneous diversification of three major clades of Gondwanan frogs at the Cretaceous-Paleogene boundary. Proc Natl Acad Sci U S A 114(29):E5864–E5870.

37. Alexander DH, Novembre J, & Lange K (2009) Fast model-based estimation of ancestry in unrelated individuals. Genome Research 19:1655–1664.

38. Pickrell JK & Pritchard JK (2012) Inference of population splits and mixtures from genome-wide allele frequency data. PLoS Genet 8(11):e1002967.

39. AmphibiaWeb (2016) Information on amphibian biology and conservation. Berkeley, California: AmphibiaWeb. Available: http://amphibiaweb.org/.

40. Hijmans RJ, Cameron, S. E., Parra, J. L., Jones, P. G. and Jarvis, A. (2005) Very high resolution interpolated climate surfaces for global land areas. Int. J. Climatol. 25:1965–1978.

41. Phillips SJ, Anderson RP, & Schapire RE (2006) Maximum entropy modeling of species geographic distributions. Ecological Modelling 190(3):231–259.

42. Novikova PY, et al. (2016) Sequencing of the genus Arabidopsis identifies a complex history of nonbifurcating speciation and abundant trans-specific polymorphism. Nat Genet 48(9):1077–1082.

43. Green RE, et al. (2010) A Draft Sequence of the Neandertal Genome. Science 328:710–722.

44. Nishihara H, Maruyama S, & Okada N (2009) Retroposon analysis and recent geological data suggest near-simultaneous divergence of the three superorders of mammals. Proc Natl Acad Sci U S A 106(13):5235–5240.

45. Hallstrom BM & Janke A (2010) Mammalian evolution may not be strictly bifurcating. Mol Biol Evol 27(12):2804–2816.

46. Garrigan D, et al. (2012) Genome sequencing reveals complex speciation in the Drosophila simulans clade. Genome Res 22(8):1499–1511.

47. Martin SH, et al. (2013) Genome-wide evidence for speciation with gene flow in Heliconius butterflies. Genome Research 23:1817–1828.

48. Jonsson H, et al. (2014) Speciation with gene flow in equids despite extensive chromosomal plasticity. Proc Natl Acad Sci U S A 111(52):18655–18660.

49. Fontaine MC, et al. (2015) Mosquito genomics. Extensive introgression in a malaria vector species complex revealed by phylogenomics. Science 347(6217):1258524.

50. Lamichhaney S, et al. (2015) Evolution of Darwin’s finches and their beaks revealed by genome sequencing. Nature 518(7539):371–375.

51. Suh A, Smeds L, & Ellegren H (2015) The Dynamics of Incomplete Lineage Sorting across the Ancient Adaptive Radiation of Neoavian Birds. PLoS Biol 13(8):e1002224.

52. Pease JB, Haak DC, Hahn MW, & Moyle LC (2016) Phylogenomics Reveals Three Sources of Adaptive Variation during a Rapid Radiation. PLoS Biol 14(2):e1002379.

53. Becak ML & Becak W (1970) Further studies on polyploid amphibians (Ceratophrydidae). 3. Meiotic aspects of the interspecific triploid hybrid: Odontophrynus cultripes (2n=22) × O. americanus (4n=44). Chromosoma 31(4):377–385.

54. Main AR (1962) Comparisons of breeding biology and isolating mechanisms in Western Australian frogs (Melbourne Univ. Press Melbourne, Victoria, Australia) pp 370–379.

55. Nishioka M & Ueda H (1983) Studies on polyploidy in Japanese frogs. Sci Rep Lab Amphibian Biol. Hiroshima Univ. 6:207–252.

56. Bogart JP & Bi K (2013) Genetic and genomic interactions of animals with different ploidy levels. Cytogenet Genome Res 140(2–4):117–136.

57. Shimada M & Hase K (2014) Female polyandry and size-assortative mating in isolated local populations of the Japanese common toad Bufo japonicus. Biological Journal of the Linnean Society 113(1):236–242.

58. Geach TJ, Stemple DL, & Zimmerman LB (2012) Genetic analysis of Xenopus tropicalis. Methods Mol Biol 917:69–110.

59. Lafon-Placette C, et al. (2017) Endosperm-based hybridization barriers explain the pattern of gene flow between Arabidopsis lyrata and Arabidopsis arenosa in Central Europe. Proc Natl Acad Sci U S A 114(6):E1027–E1035.

60. Lafon-Placette C & Kohler C (2016) Endosperm-based postzygotic hybridization barriers: developmental mechanisms and evolutionary drivers. Mol Ecol 25(11):2620–2629.

61. Schmickl R, Marburger S, Bray S, & Yant L (2017) Hybrids and horizontal transfer: introgression allows adaptive allele discovery. J Exp Bot 68(20):5453–5470.

62. McCartney-Melstad E & Shaffer HB (2015) Amphibian molecular ecology and how it has informed conservation. Mol Ecol 24(20):5084–5109.

63. McCartney-Melstad E, Gidis M, & Shaffer HB (2018) Population genomic data reveal extreme geographic subdivision and novel conservation actions for the declining foothill yellow-legged frog. Heredity (Edinb) 121(2):112–125.

64. Callaghan CT, Rowley JJL, Cornwell WK, Poore AGB, & Major RE (2019) Improving big citizen science data: Moving beyond haphazard sampling. PLoS Biol 17(6):e3000357.

65. Rowley JJL, et al. (2019) FrogID: Citizen scientists provide validated biodiversity data on frogs of Australia. Herpetological Conservation and Biology 14(1):155–170.

66. Prum RO, et al. (2015) A comprehensive phylogeny of birds (Aves) using targeted next-generation DNA sequencing. Nature 526:569.

67. Rokyta DR, Lemmon AR, Margres MJ, & Aronow K (2012) The venom-gland transcriptome of the eastern diamondback rattlesnake (Crotalus adamanteus). BMC Genomics 13:312.

68. Hamilton CA, Lemmon AR, Lemmon EM, & Bond JE (2016) Expanding anchored hybrid enrichment to resolve both deep and shallow relationships within the spider tree of life. BMC Evol Biol 16(1):212.

69. Pyron RA, Hsieh FW, Lemmon AR, Lemmon EM, & Hendry CR (2016) Integrating phylogenomic and morphological data to assess candidate species-delimitation models in brown and red-bellied snakes (Storeria). Zoological Journal of the Linnean Society 177(4):937–949.

70. Katoh K & Standley DM (2013) MAFFT multiple sequence alignment software version 7: improvements in performance and usability. Mol Biol Evol 30(4):772–780.

71. Kearse M, et al. (2012) Geneious Basic: an integrated and extendable desktop software platform for the organization and analysis of sequence data. Bioinformatics 28(12):1647–1649.

72. Taucce PPG, et al. (2018) The mitochondrial genomes of five frog species of the Neotropical genus Ischnocnema (Anura: Brachycephaloidea: Brachycephalidae). Mitochondrial DNA Part B 3(2):915–917.

73. Irisarri I, et al. (2012) The origin of modern frogs (Neobatrachia) was accompanied by acceleration in mitochondrial and nuclear substitution rates. BMC Genomics 13(1):626.

74. Hahn C, Bachmann L, & Chevreux B (2013) Reconstructing mitochondrial genomes directly from genomic next-generation sequencing reads—a baiting and iterative mapping approach. Nucleic Acids Research 41(13):e129–e129.

75. Edgar RC (2004) MUSCLE: multiple sequence alignment with high accuracy and high throughput. Nucleic Acids Res 32(5):1792–1797.

76. Hillis DM, Heath TA, & St John K (2005) Analysis and visualization of tree space. Syst Biol 54(3):471–482.

77. Paradis E, Claude J, & Strimmer K (2004) APE: Analyses of Phylogenetics and Evolution in R language. Bioinformatics (Oxford, England) 20:289–290.

78. Maechler M, Rousseeuw P, Struyf A, & Hubert M (2018) cluster: Cluster Analysis Basics and Extension.

79. Weiss CL, Pais M, Cano LM, Kamoun S, & Burbano HA (2018) nQuire: a statistical framework for ploidy estimation using next generation sequencing. BMC Bioinformatics 19(1):122.

80. Li H & Durbin R (2009) Fast and accurate short read alignment with Burrows-Wheeler transform. Bioinformatics (Oxford, England) 25:1754–1760.

81. Li H, et al. (2009) The Sequence Alignment/Map format and SAMtools. Bioinformatics (Oxford, England) 25:2078–2079.

82. Roberts JD (1997) Geographic Variation in Calls of Males and Determination of Species Boundaries in Tetraploid Frogs of the Australian Genus Neobatrachus (Myobatrachidae). 45(2):95–112.

83. Mahony M (1986) Cytogenetic studies on Australian frogs of the family Myobatrachidae. Ph.D. thesis, Macquarie University, Sydney, Australia.

84. Choleva L & Janko K (2013) Rise and persistence of animal polyploidy: evolutionary constraints and potential. Cytogenet Genome Res 140(2–4):151–170.

85. Okamoto T, Ohnishi Y, & Toda E (2017) Development of polyspermic zygote and possible contribution of polyspermy to polyploid formation in angiosperms. J Plant Res 130(3):485–490.

86. Hollister JD, et al. (2012) Genetic adaptation associated with genome-doubling in autotetraploid Arabidopsis arenosa. PLoS Genet 8(12):e1003093.

87. Arnold B, Kim ST, & Bomblies K (2015) Single Geographic Origin of a Widespread Autotetraploid Arabidopsis arenosa Lineage Followed by Interploidy Admixture. Mol Biol Evol 32(6):1382–1395.

88. Ramsey J & Schemske DW (2002) Neopolyploidy in Flowering Plants. Annual Review of Ecology and Systematics 33(1):589–639.

89. Pfeifer B, Wittelsburger U, Ramos-Onsins SE, & Lercher MJ (2014) PopGenome: an efficient Swiss army knife for population genomic analyses in R. Mol Biol Evol 31(7):1929–1936.

90. Hijmans RJ, van Etten J. (2012) raster: Geographic analysis and modeling with raster data. R package version 2.0-12.

