## Supplementary Material for "Whole genome duplication potentiates inter-specific hybridisation and niche shifts in Australian burrowing frogs *Neobatrachus*"

**Appendix 1. Summary of cytogenetic observations and mechanisms for uni-directional introgression**

Evidence for unreduced gamete formation

Cytogenetic evidence

Diploid *Neobatrachus* can produce unreduced gametes, as evidenced cytogenetically by a triploid *fulvus* (2n) x *sutor* (2n) hybrid (Fig. S12C).

Evidence for triploid formation comes from three sources:

Cytogenetic evidence

A triploid *fulvus* (2n) x *sutor* (2n) hybrid (Fig. S12C).

Triploid *pictus* (2n) x *sudellae* (4n) hybrids (Fig. S12B) have been observed at hybrid zones between *pictus* and *sudellae* on the Eyre Peninsula, South Australia and at Moyston east of the Grampians, Victoria (Mahony 1986, unpublished data) (Table S3).

Lab crosses

Evidence of possible triploid formation comes from another diploid x tetraploid species combination from viable laboratory crosses of *N. sutor* (2n) x *N. kunapalari* (4n) produced by Main (1962) (Table S3).

Molecular genetic evidence

Evidence of triploid formation through hybridisation comes from our inference of a triploid resulting from a cross between *wilsmorei* (2n) x *kunapalari* (4n) or *sudellae* (4n) from the exon analysis of WAM R119302 from Yalgoo, WA [see Supplementary Data on ID corrections].

“Triploid Bridge”

Triploids can produce 3n gametes, as evidenced by the presence of a pentaploid *pictus* x *sudellae* hybrid (Fig. S12F) at a *pictus* (2n) x *sudellae* (4n) hybrid zone at Moyston, east of the Grampians, Victoria where triploid *pictus* x *sudellae* hybrids were observed also (Mahony 1986, unpublished data) (Table S3).

A triploid producing an unreduced gamete (3n) in a cross with a diploid would produce a tetraploid [no direct observation so far] which could backcross to a 4n individual producing introgression from the diploid into the tetraploid gene pool, i.e the so-called “triploid bridge”.

Alternatively triploid frogs can in some cases produce balanced haploid, diploid or triploid gametes (Becak and Becak 1970) which then opens the possibility of crosses with diploids or tetraploid that would produce tetraploids that could backcross into the sympatric tetraploid gene pool.

Other mechanisms of diploid to tetraploid gene flow

A diploid could produce an unreduced gamete and cross with a tetraploid to produce a novel tetraploid that could backcross into the parental tetraploid population. While we don’t have unequivocal direct observations of this so far, the tetraploid hybrids (Table S3) are either of a *kunapalari* x *sudellae* or a *kunapalari* x diploid spp. origin (Fig. S12C-E). The karyotypes of *sudellae* and some diploids that are sympatric with *kunapalari* do not have diagnostic chromosomal markers (Roberts et al. 1991) and as tissue samples from these individuals are not available we presently are not able to distinguish these alternative scenarios.

Early embryos can be experimentally shocked into suppressing cleavage divisions to produce an autopolyploid (Nishioka & Ueda 1983), which could then mate with sympatric tetraploids. Cold or heat shocks are possible for *Neobatrachus* eggs clutches at numerous locations across southern Australia. For instance Roberts and Edwards (2018) note that the minimum maximum temperature range for Hyden WA as −5.6 °C to 48.6 °C. Breeding by more southerly *Neobatrachus* tends to follow the onset of autumn and winter rains, typical of the Mediterranean climate in this region. Exposure to freezing or near freezing conditions in shallow egg deposition sites following cold fronts associated with rain fall events is frequently possible. *Neobatrachus* in the arid zone breed in association with heavy summer rains which may be associated with heat wave conditions.

Egg production by hybrid females could have an inflated likelihood of the production of unreduced gametes or hybrid zygotes could have an elevated risk of the suppression of the first mitosis following fertilization.

**References**

Becak, M. L., Becak, W. 1970. Further studies on polyploid amphibians (Ceratophrydidae). 111. Meiotic aspects of the interspecific triploid hybrid: *Odontophrynus cultripes* (2n = 22) *x 0. americanus* (4n = 44). *Chromosoma* **31**, 377-85.

Main, A. R. 1962. Comparisons of breeding biology and isolating mechanisms in Western Australian frogs, p. 370-379. In: *The evolution of living organisms*. G. W. Leeper (ed.). Melbourne Univ. Press, Melbourne, Victoria, Australia.

Nishioka M, Ueda H 1983.Studies on polyploidy in Japanese frogs. *Sci Rep Lab Amphibian Biol. Hiroshima Univ*. **,**: 207–252.

Roberts, JD., Edwards, D. 2018. The Evolution, Physiology and Ecology of the Australian Arid-Zone Frog Fauna, p. 149-180. In: *On the Ecology of Australia’s Arid Zone*. Springer, Cham

Roberts, J. D., M. J. Mahony, P. Kendrick, and C. M. Majors. 1991. A new species of burrowing frog, *Neobatrachus* (Anura: Myobatrachidae), from the eastern wheatbelt of Western Australia. *Records of the Western Australian Museum* **15**, 23–32.

**Supplementary Table 1.** Summary statistics for each of the species calculated with R package “PopGenome”

| Species | Number of samples | Pi median, % | CI_pi_median, % | Tajima's *D* median | Confidence interval Tajima's *D* median | Number of valid loci | Average number of valid sites in loci |
| --- | --- | --- | --- | --- | --- | --- | --- |
| *N. albipes* | 8 | 0.161 | [0.1398, 0.184] | -0.853 | [-0.9717, -0.7762] | 415 | 1422 |
| *N. aquilonius* | 11 | 0.661 | [0.5021, 0.9544] | -1.309 | [-1.416, -1.207] | 233 | 643 |
| *N. fulvus* | 8 | 0.127 | [0.09706, 0.165] | -0.955 | [-1.055, -0.7119] | 224 | 1192 |
| *N. kunapalari* | 8 | 1.638 | [1.379, 2.102] | -0.509 | [-0.57, -0.4297] | 249 | 741 |
| *N. pelobatoides* | 7 | 0.226 | [0.2022, 0.2594] | -0.956 | [-1.075, -0.8798] | 331 | 1000 |
| *N. pictus* | 5 | 0.296 | [0.2655, 0.3195] | -0.11 | [-0.1885, -0.02694] | 408 | 1450 |
| *N. sudellae* | 14 | 1.248 | [0.9262, 1.947] | -1.404 | [-1.474, -1.347] | 215 | 575 |
| *N. sutor* | 9 | 0.288 | [0.2658, 0.3358] | -0.533 | [-0.6178, -0.3964] | 222 | 1016 |
| *N. wilsmorei* | 5 | 0.114 | [0.1059, 0.1294] | -0.508 | [-0.626, -0.3559] | 421 | 1388 |

**Supplementary Table 2.** The average test AUC (area under the Receiving Operator Curve) for the replicate runs for all the species in MaxEnt modeling for predicting species distribution from climate data at the species occurrences.

| species | AUC | AUC standard deviation |
| --- | --- | --- |
| *N. albipes* | 0,988 | 0,003 |
| *N. aquilonius* | 0,923 | 0,011 |
| *N. fulvus* | 0,999 | 0 |
| *N. kunapalari* | 0,964 | 0,004 |
| *N. pelobatoides* | 0,988 | 0,002 |
| *N. pictus* | 0,977 | 0,004 |
| *N. sudellae* | 0,908 | 0,008 |
| *N. sutor* | 0,944 | 0,006 |
| *N. wilsmorei* | 0,978 | 0,004 |

**Supplementary Table 3.** Instances of polyploid *Neobatrachus*

| id_num | Sex | Putative parental taxa | | Ploidy | Life stage | Locality | State | Reference |
| --- | --- | --- | --- | --- | --- | --- | --- | --- |
|  | F | fulvus 2n | sutor 2n | 3n | adult | Learmonth | WA | Mahony 1986 |
|  |  | pictus 2n | sudellae 4n | 3n | adult | Eyre Peninsula | SA | Mahony 1986 |
|  | - | pictus 2n | sudellae 4n | 3n | metamorph | Moyston, east of Grampians | Vic | Mahony 1986 |
|  | - | pictus 2n | sudellae 4n | 5n | metamorph | Moyston, east of Grampians | Vic | Mahony unpubl. data |
|  |  | kunapalari 4n | sudellae 4n | 4n | adult | North of Menzies | WA | Mahony 1986 |
|  | | kunapalari 4n | unknown 4n | 4n | adult | Ninghan | WA | Mahony 1986 |
| I5442/WAMR119302 | | wilsmorei 2n | kunapalari or sudellae 4n | 3n |  | Yalgoo | WA | this paper |
|  |  | sutor 2n | kunapalari 4n | 3n | - | Lab cross | WA | Main 1962 |

**Fig. S1.** Species tree and admixture results for optimal clustering at K equals 3, 7 and 9. Vertical colored bars to the left of the tips of the tree correspond to our final species assignments (Supplementary Table 1); colors of the bars are species-specific and correspond to the branch colors from Fig. 1A; filtered out samples are marked with black bars.

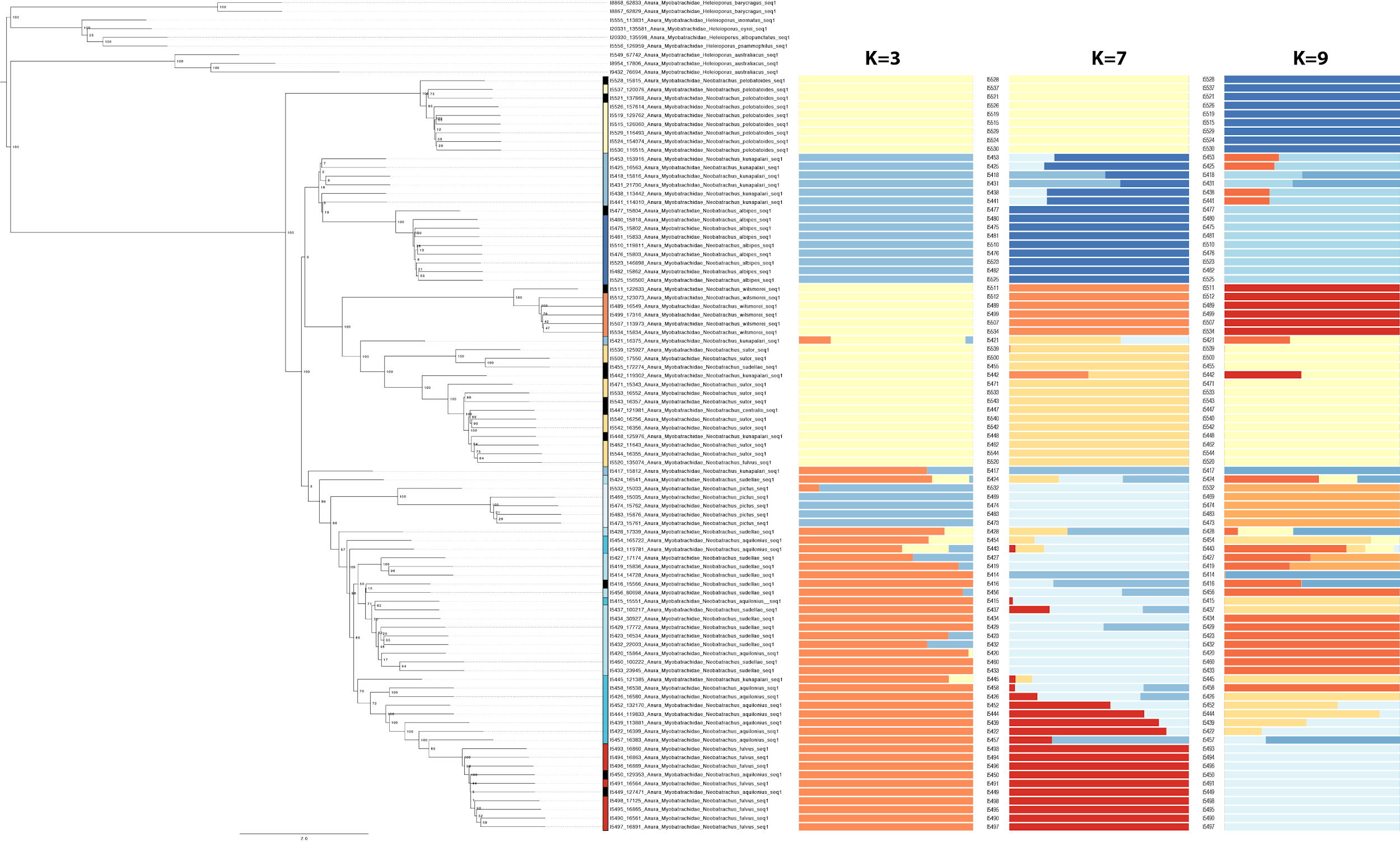

**Fig. S2**. Nuclear species tree as inferred using ASTRAL, all nuclear loci, and complete taxon sampling.
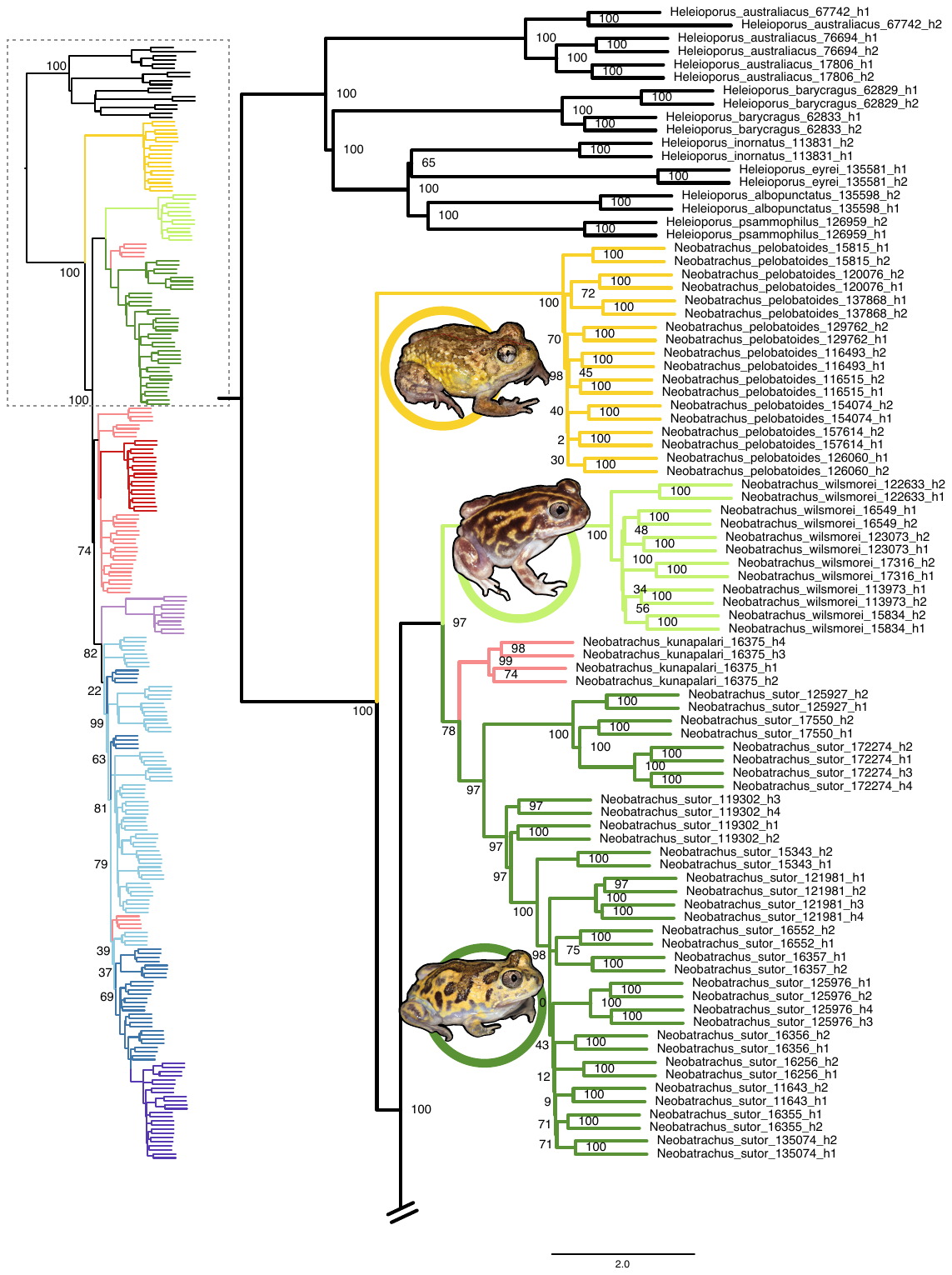

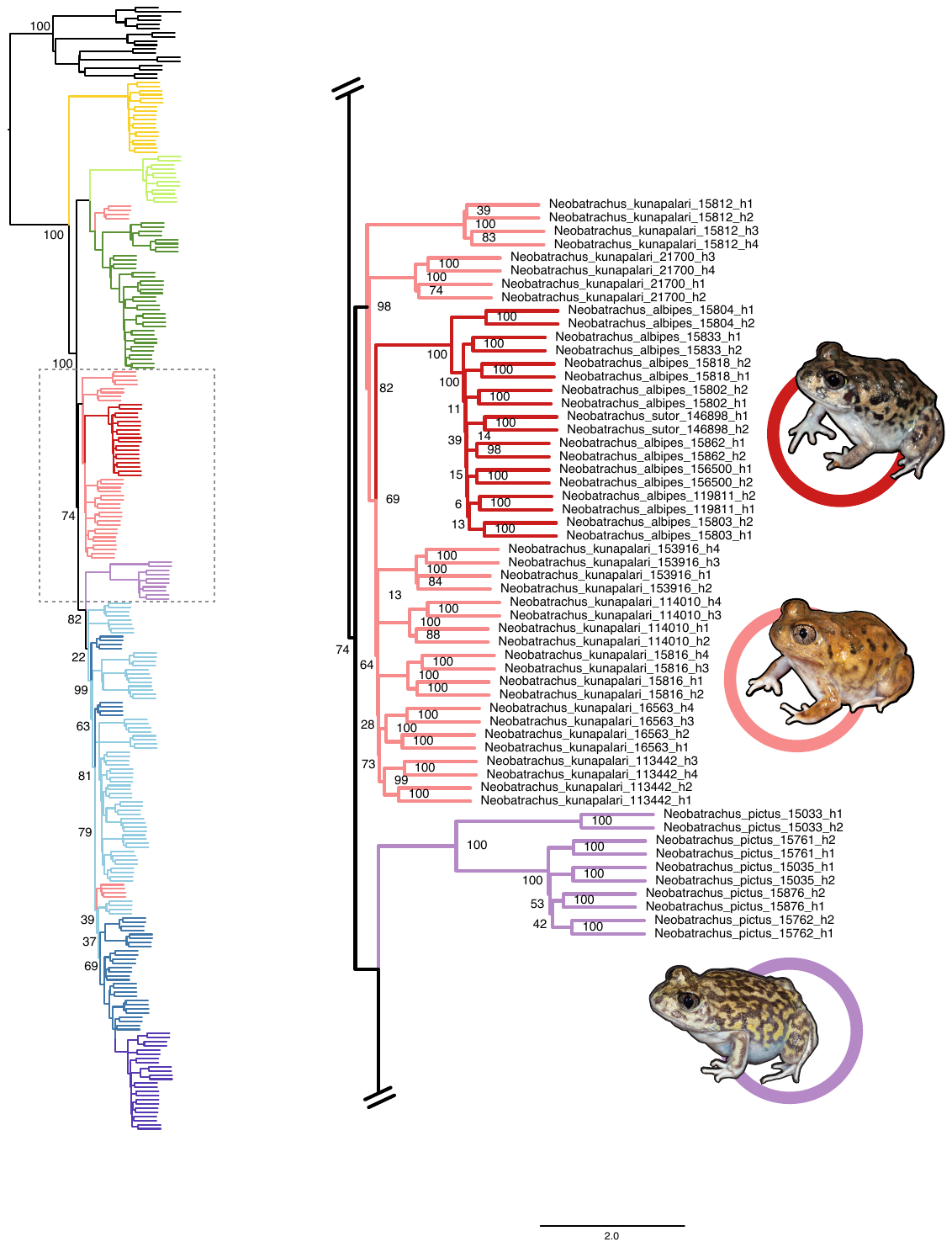

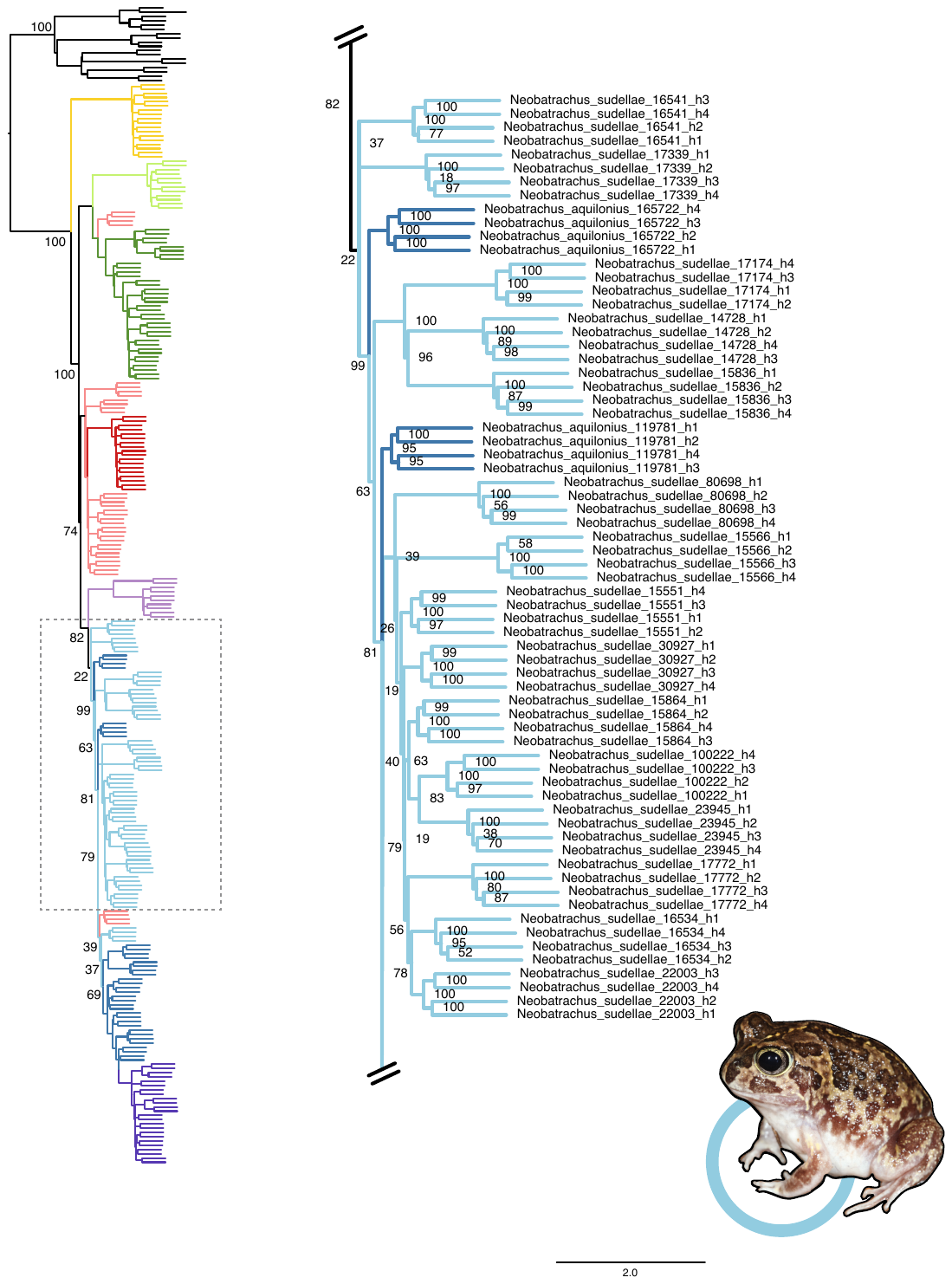

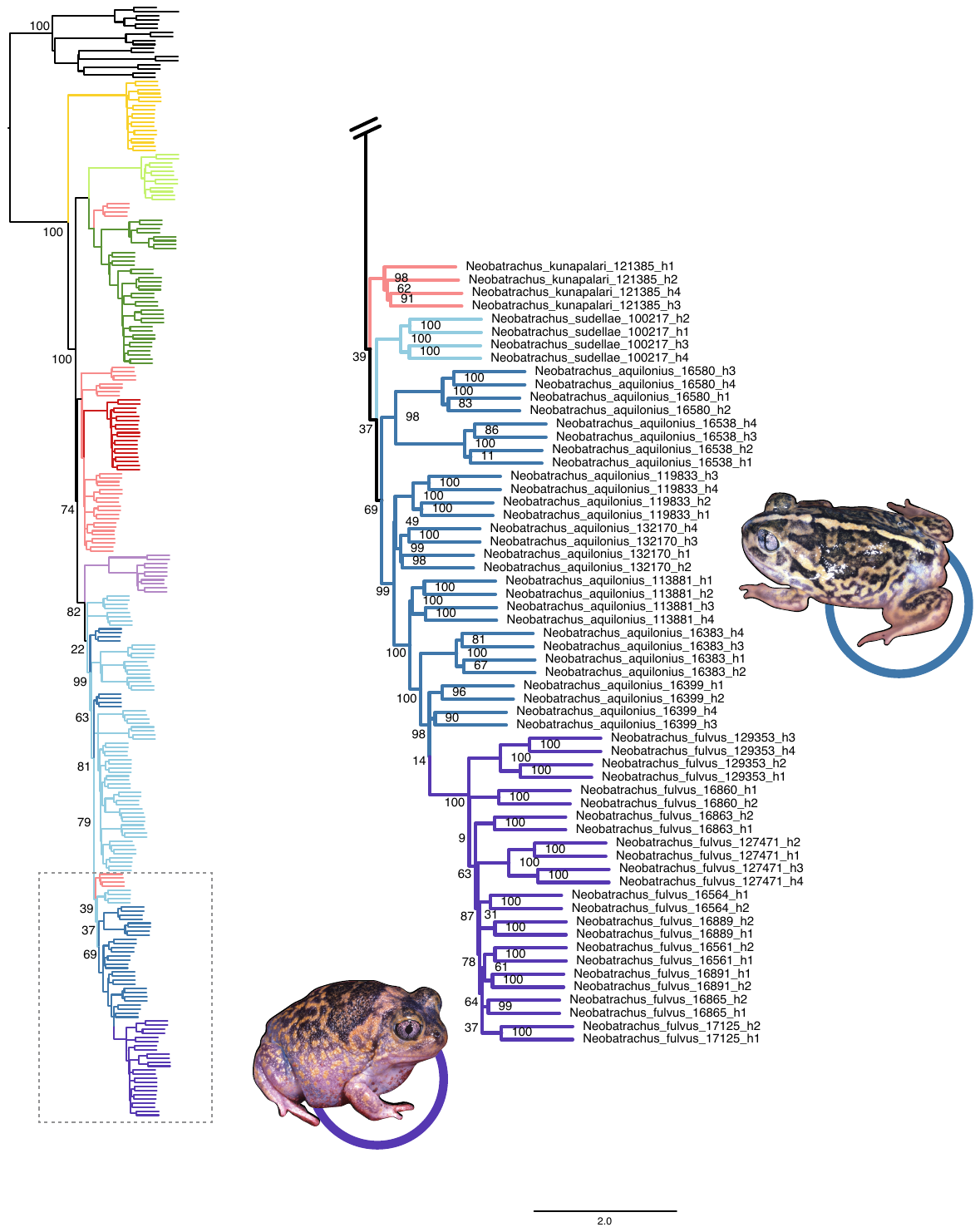

**Fig. S3.** Two dimensional representations of MDS gene tree space, colored by optimal clustering scheme for two dimensions (k=2) and three dimensions (k=4), and their associated topologies inferred using ASTRAL. Each point represents a single gene tree, colored clusters match colored trees displayed to the right. Nodes at values indicate bootstrap support.

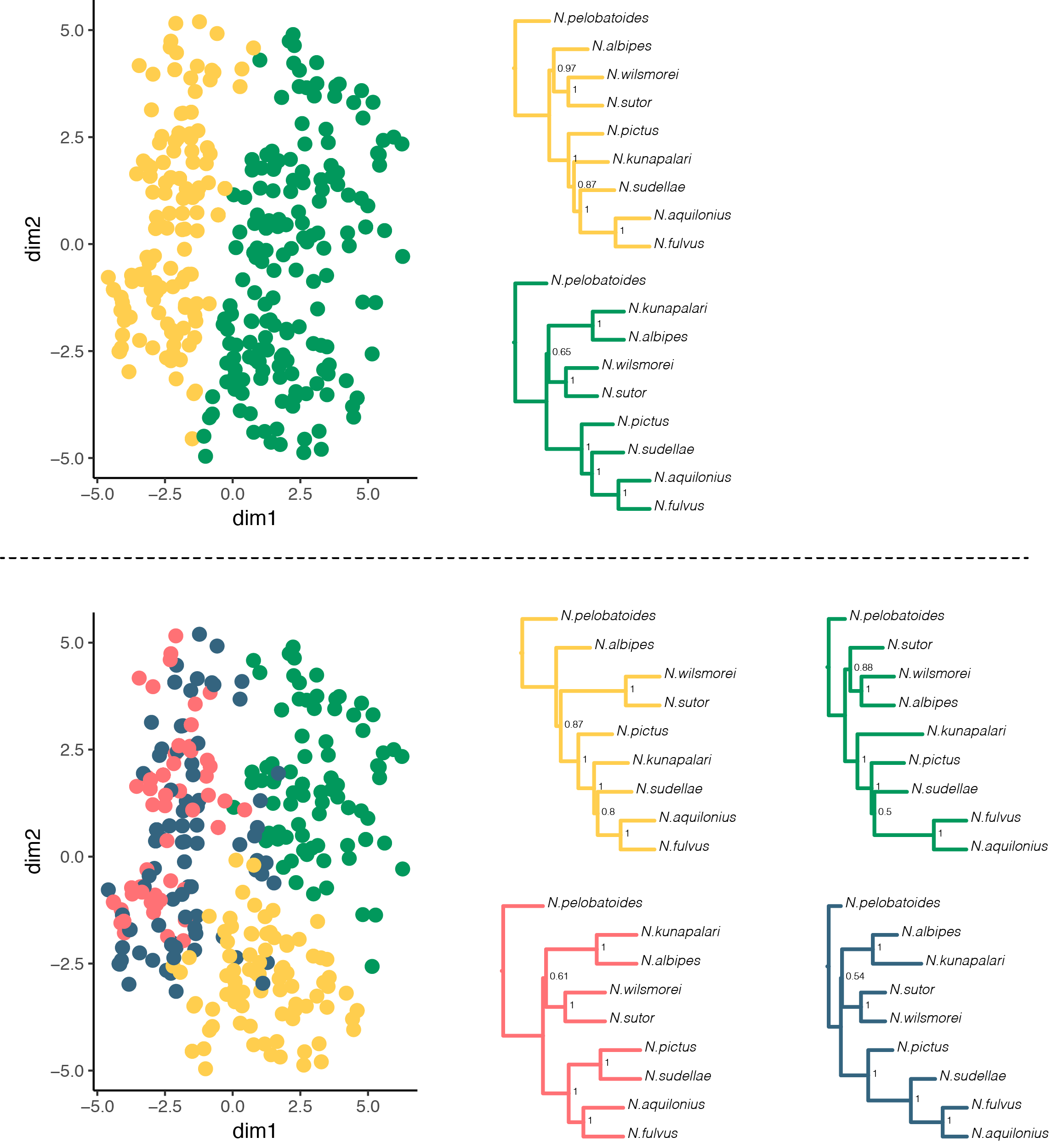

**Fig. S4.** Distribution of allele frequencies of biallelic sites in *Neobatrachus* tetraploids supports tetrasomic inheritance mode in *N. sudellae* and *N. aquilonius* and mixed inheritance mode in *N. kunapalari*. (A) Pairwise combination of individuals within the diploid species model the expected allele frequencies in autotetraploids with tetrasomic inheritance (blue line), when pairwise combination of individuals between the diploid *Neobatrachus* species model the expected distribution for allotetraploids with disomic inheritance mode (purple line). Modeled allotetraploids show excess of intermediate allele frequencies compared to autotetraploids. Gray area shows 95% confidence interval. (B) Comparing the ratio between intermediate (40-60%) and rare (<30%) allele frequencies we reject allotetraploid origin for *N. sudellae* and *N. aquilonius*, when *N. kunapalari* shows intermediate distribution, suggesting mixed inheritance. Comparisons performed with Wilcoxon tests adjusted for multiple testing.

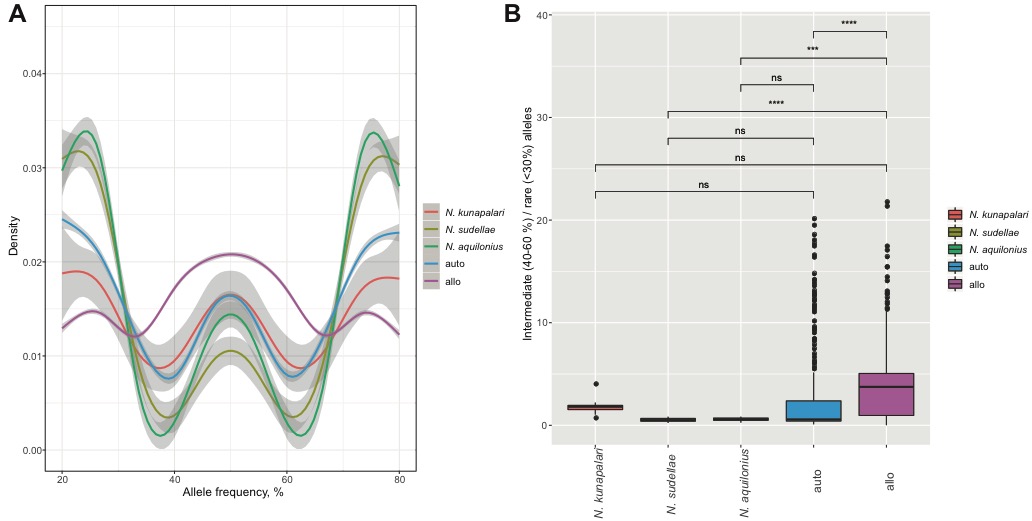

**Fig.S5.** Cross-validation plot showing three local optimal solutions for ADMIXTURE clustering at K equals 3, 7 and 9.

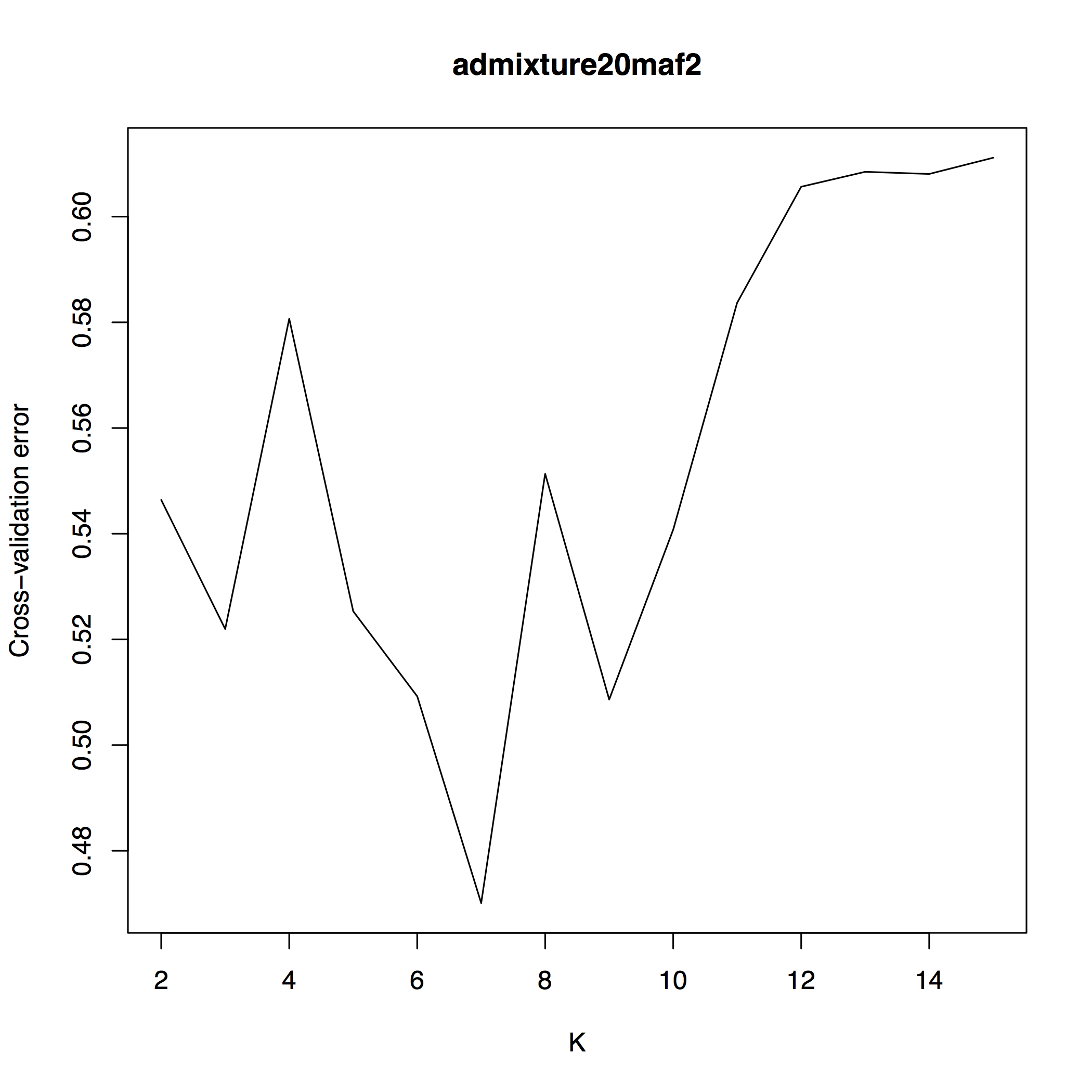

**Fig. S6.** Heatmap and hierarchical clustering of the *Neobatrachus* lineages based on the distance matrix from pairwise median Fst values. Tetraploid species (*N. sudellae*, *N. aquilonius* and *N. kunapalari*; highlighted with black left bar) cluster together and are characterised by the lowest Fst values between each other. This, together with low Fst values between tetraploid and diploid lineages, can probably be explained by the gene flow within the tetraploids and between the diploids and the tetraploids. Diploid lineages (highlighted with grey left bar) appear to be more isolated from each other compared to tetraploids, which is in agreement with ADMIXTURE assignment results and TreeMix estimations of possible migration events.

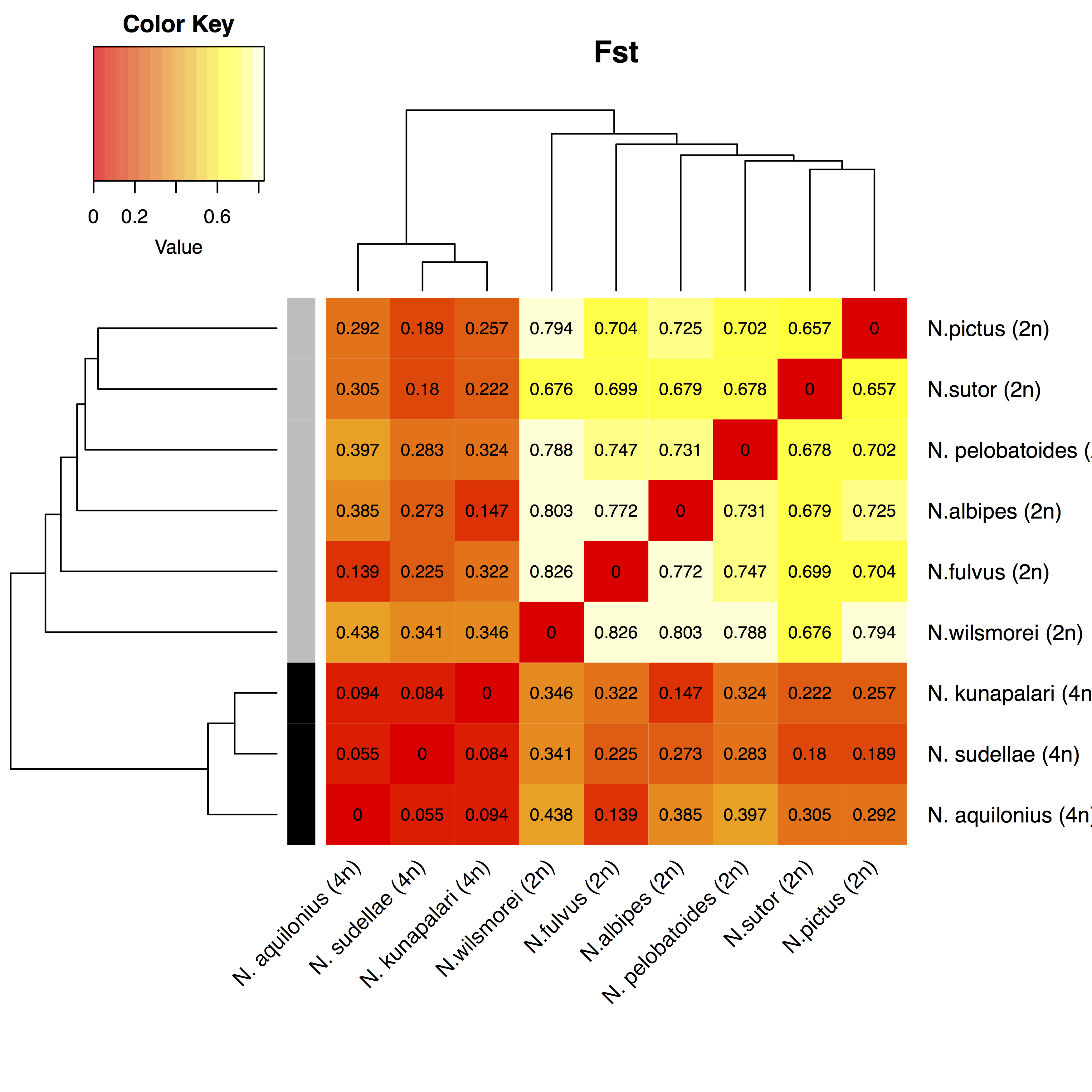

**Fig.S7**. Occurrence data locations registered at the AmphibiaWeb database for *Neobatrachus* species: A - tetraploids, B - diploids.

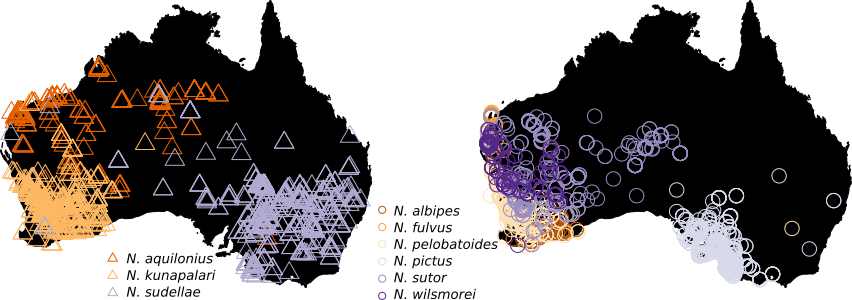

**Fig. S8.** PCA analysis of bioclimatic variables for *Neobatrachus* entries in the occurrence AmphibiaWeb database. A) Barplot showing the percentage of variances explained by each principal component. The first three principal components are labeled with the top three contributions of variables. BIO10 = Mean Temperature of Warmest Quarter, BIO12 = Annual Precipitation, BIO17 = Precipitation of Driest Quarter, BIO18 = Precipitation of Warmest Quarter, BIO19 = Precipitation of Coldest Quarter. B-D) Pairwise combinations of the first three principal components, where individuals with a similar profile of bioclimatic data are grouped together. Points represent each individual and colored according to the species assignment, ellipses represent 95% confidence area.

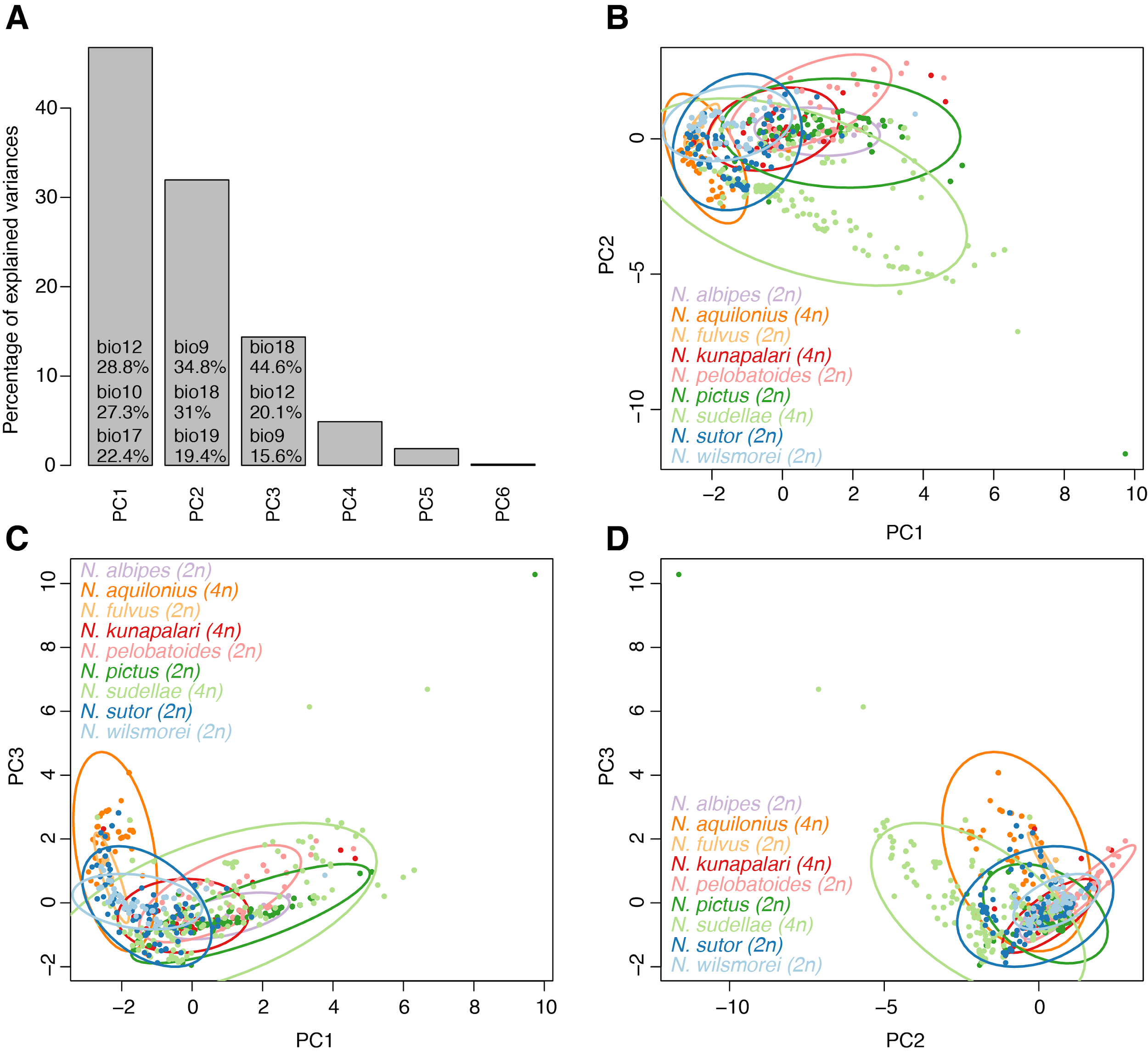

**Fig. S9.** The results of the jackknife test of variable importance for models on each species. BIO19 (Precipitation of Coldest Quarter) was the most informative variable for the models of *N. pelobatoides* and *N. albipes* distributions; BIO18 (Precipitation of Warmest Quarter) was the most informative variable for the models of *N. wilsmorei*, *N. sutor* and *N. kunapalari*; BIO17 (Precipitation of Driest Quarter) was the most informative variable for the model of *N. fulvus*; BIO10 (Mean Temperature of Warmest Quarter) for *N. pictus*; and BIO9 (Mean Temperature of Driest Quarter) for *N. sudellae* and *N. aquilonius*.

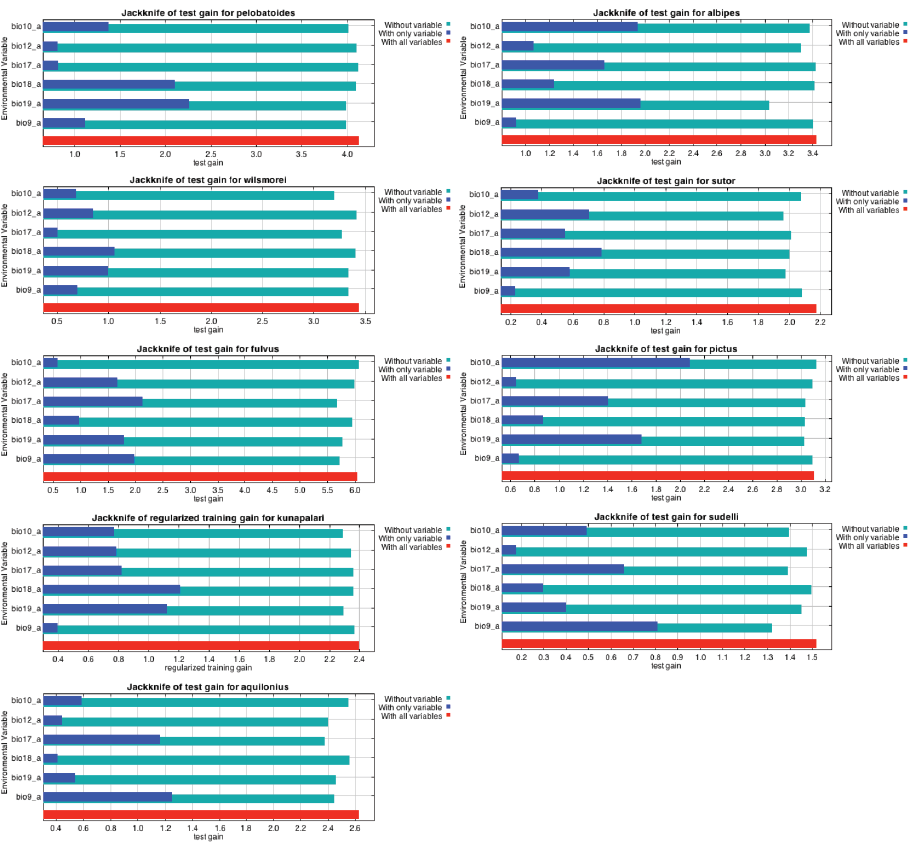

**Fig. S10.** The point-wise mean of the 10 models for each of the diploid species build on environmental layers from the current climate data and applied to the environmental layers from the Last Glacial Maximum climate data.

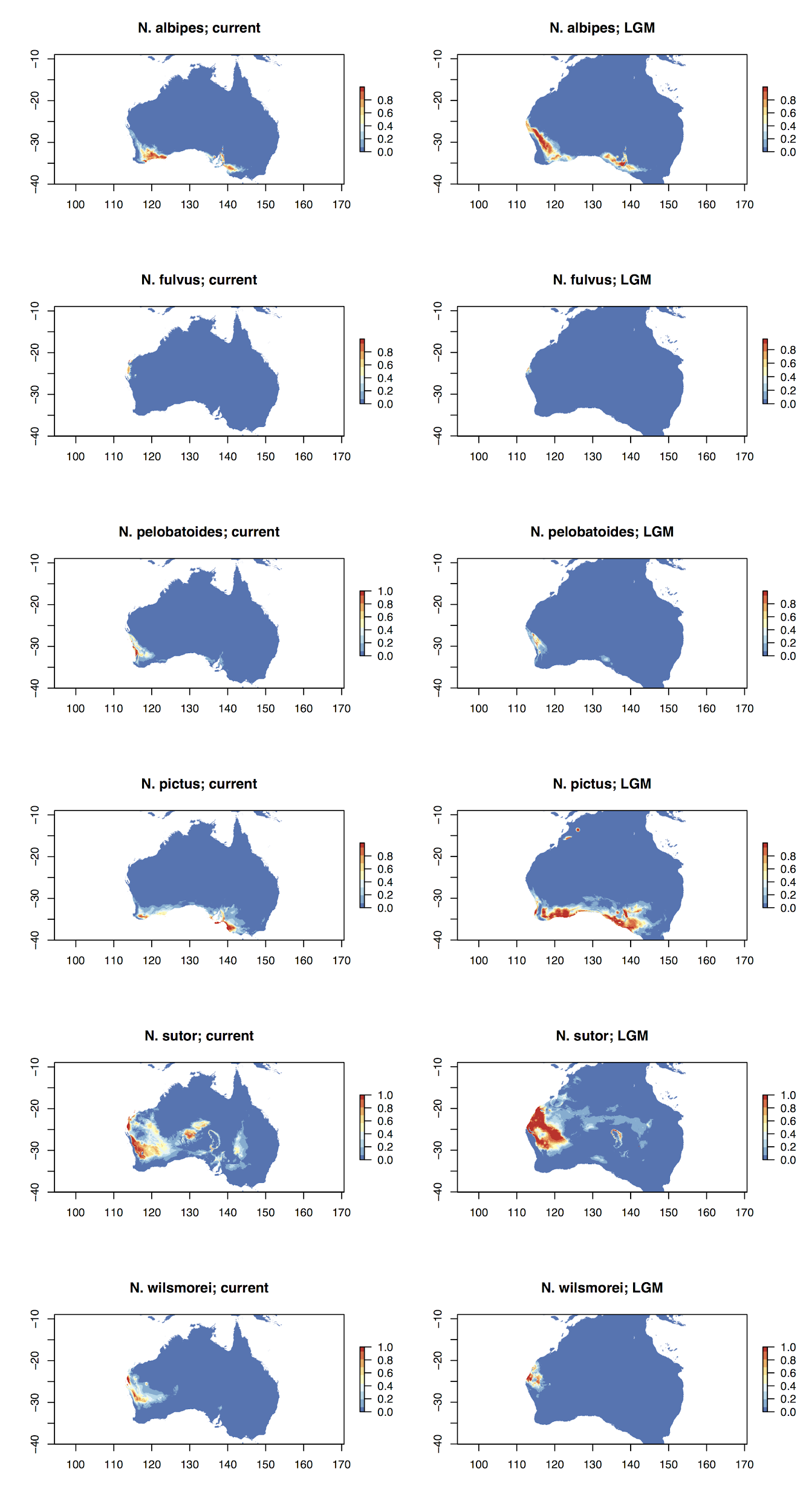

**Fig. S11**. The point-wise mean of the 10 models for each of the tetraploid species build on environmental layers from the current climate data and applied to the environmental layers from the Last Glacial Maximum climate data.

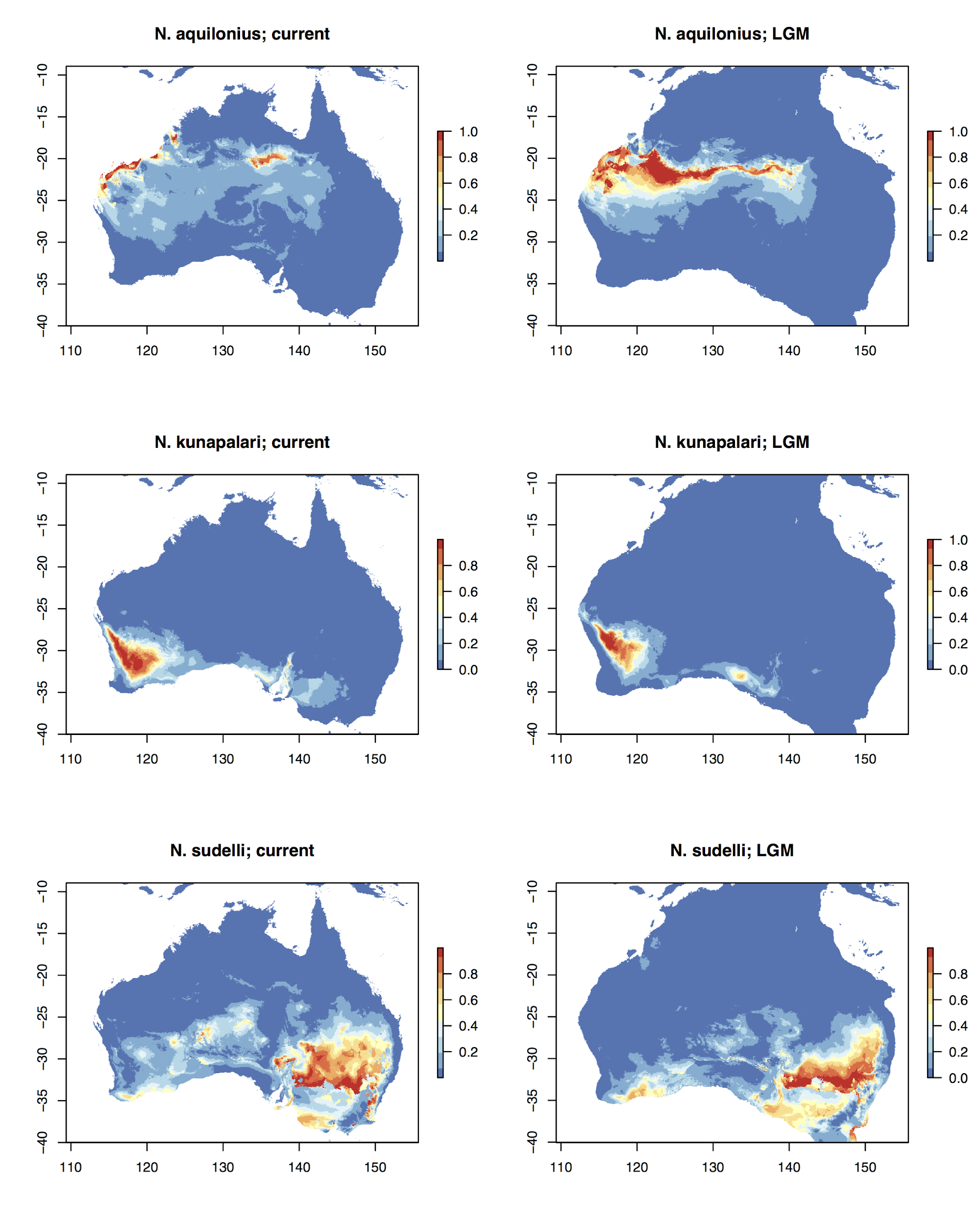

**Fig. S12.** Karyotypes of *Neobatrachus*. **A**) *N. sutor* [2n], **B**) *N. pictus* x *N. sudellae* triploid [3n] hybrid from Moyston, east of the Grampians, Victoria, **C**) *N. fulvus* x *N. sutor* triploid [3n] hybrid from Learmonth, Western Australia, **D**) *N. sudellae* [4n], **E**) tetraploid x tetraploid hybrid from north of Menzies, Western Australia, **F**) *N. pictus* x *N. sudellae* pentaploid [5n] hybrid from Moyston, east of the Grampians, Victoria. Arrowheads indicate nucleolar organiser regions (NORs).

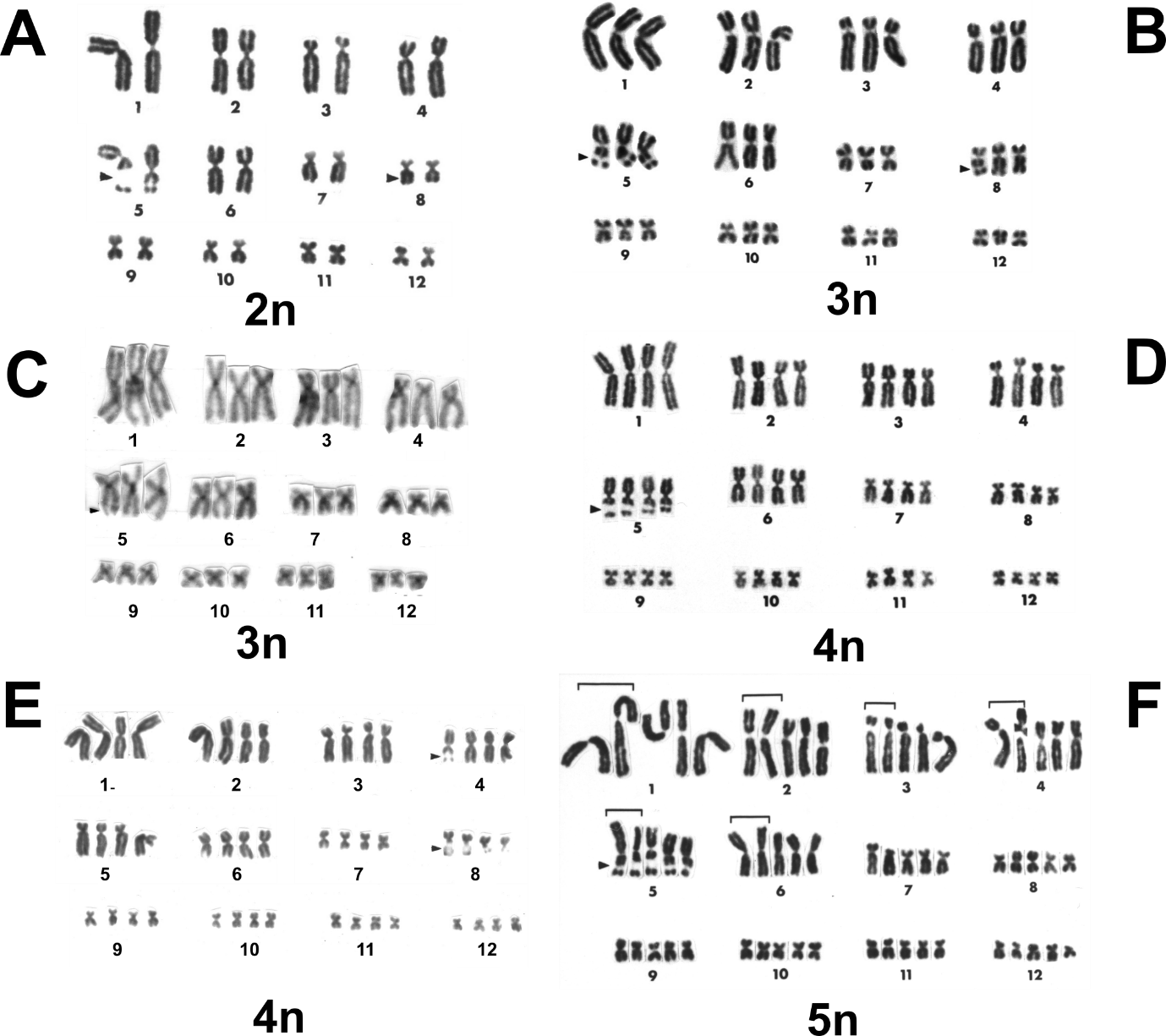
